# Mechanistic insights into the color transformation of a non-FRET substrate for RNase activity detection

**DOI:** 10.64898/2026.03.11.710959

**Authors:** Sohyun Kim, Soonwoo Hong, Jada N. Walker, Yujie He, Anh-Thu Nguyen, Wei-Ru Chen, Saeed Seifi, Yuan-I Chen, Yu-An Kuo, Jennifer S. Brodbelt, Hsin-Chih Yeh

## Abstract

DNA-templated silver nanoclusters (DNA/AgNCs) have created a new class of non-FRET DNase substrates, termed Subak, that exhibits a color change upon DNase digestion. Although Subak substrates offer advantages such as ratiometric readouts and low manufacturing costs over traditional FRET substrates, the mechanism governing AgNC color switching remains unclear. Here, using a site-specific cleavage strategy, we identify color-switching hotspots and demonstrate that AgNC transformation can be controlled by the cleavage positions within the nucleic acid host. Our data support a cleavage-driven reorganization of the AgNC coordination environment, converting a non-emissive precursor into a red-emitting cluster, rather than direct enzyme-cluster interactions. Leveraging this insight, we engineer rSubak, an RNA-incorporated Subak that displays 95 nm red shift (530 to 625 nm) upon RNase cleavage. In amplification-free CRISPR/Cas13 assays for SARS-CoV-2, influenza A (A/H5N1), and measles viruses (MV) detection, rSubak achieved a limit of detection of 0.3 pM, superior to that of the commercial RNaseAlert (∼250 pM). Collectively, our results establish Subak as a generalizable, non-FRET platform for sensitive ratiometric reporting the activities of diverse nucleases.

## INTRODUCTION

Interest in using non-FRET (Förster Resonance Energy Transfer) probes in bioimaging and diagnostics has grown rapidly in recent years^1^. Because the fluorescence properties of these probes are highly sensitive to their local microenvironment, molecular binding and conformational changes can modulate their fluorescence signatures. For instance, circularly permuted GFP (cpGFP) exhibits enhanced fluorescence upon conformational rearrangements around the chromophore^2^, enabling the development of small molecule- and metal ion-sensing probes such as dLight^3^ and GCaMP^4^. Similarly, Janelia Fluor (JF) dyes conjugated to HaloTag display changes in fluorescence intensity and lifetime^5^ in response to changes in the dyes’ microenvironment^6,7^. Furthermore, fluorogenic aptamers provide a complementary strategy in which aptamer mutations tune the emission spectrum or lifetime of bound fluorogens such as DFHO and TO1-biotin^8,9^. Unlike FRET probes, these non-FRET probes eliminate donor-acceptor pairs and bypass constraints associated with donor-acceptor separation distance, resulting in smaller probe sizes and reduced manufacturing costs^1,2^. However, most non-FRET probes are not designed as cleavable enzyme substrates and typically do not provide ratiometric readouts due to the lack of a substantial emission-color shift upon activation^8^. Consequently, a low-cost non-FRET substrate that exhibits a large color change upon enzymatic cleavage would be highly distinctive and particularly valuable for molecular assays currently dominated by FRET substrates.

Based on DNA-templated silver nanoclusters (DNA/AgNCs)^10–14^, we have previously created Subak, a class of low-cost, non-FRET substrates that exhibits remarkable green-to-red color switching upon DNase cleavage^1^ (with an emission peak shift of 90-95 nm, **Fig. 1**). Compared to the commonly used commercial DNase substrate, DNaseAlert^15^, Subak offers ratiometric readouts in the clustered regularly interspaced short palindromic repeats (CRISPR)-based assays and achieving a broader sensing dynamic range at only a fraction of the cost. Although transformations of gold or silver nanoclusters from one species to another^16,17^ (e.g., Au_25_(SR)_18_ to Au_38_(SR)_24_) or establishments of dynamic equilibria among various cluster species^18–21^ (e.g., Ag_14_^8+^ ↔ Ag_11_^7+^ and Ag_20_^10+^ ↔ Ag_27_^17+^ ↔ Ag_34_^24+^) have been reported, none of these transformations have been exploited as enzyme substrates that fluoresce differently upon enzymatic processing, underscoring the uniqueness of Subak. While Subak outlines a new strategy for creating non-FRET probes for molecular biology assays, its underlying mechanisms and transformation pathways remain poorly understood. One proposed mechanism for the observed green-to-red color conversion upon DNase digestion is cluster fragmentation, which reduces the number of neutral Ag atoms in the cluster core. However, alternative transformation pathways, including redox reaction^20^, cluster fusion^17^, atom exchange^19^ and geometry change (proposed in this report), which can alter either the valence electron count or the electronic structure of the silver cluster core, should not be excluded.

**Fig 1:**
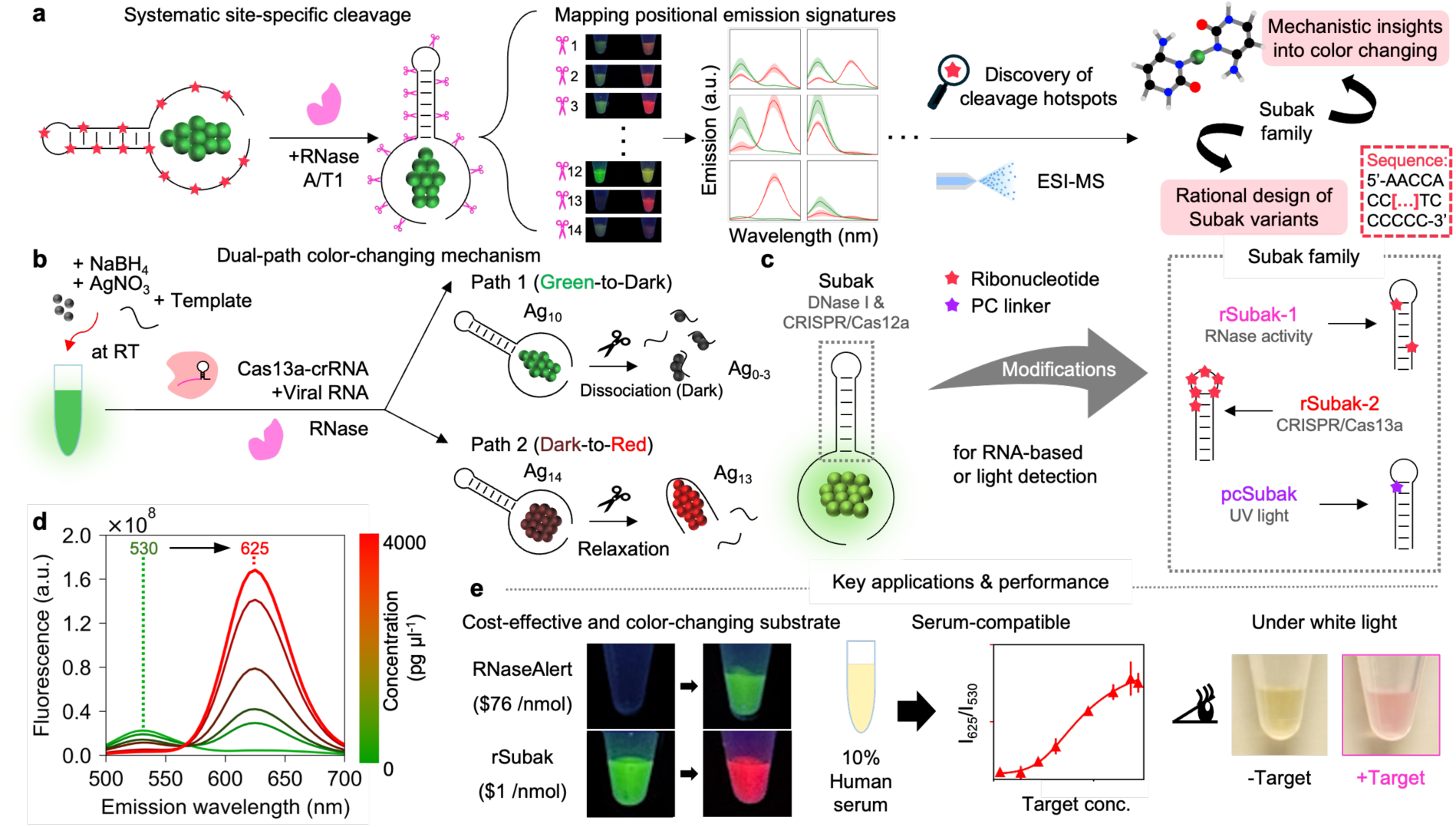
Design, mechanism, and performance of non-FRET Subak substrate. **a,** By systematically substituting selected deoxyribonucleotides with ribonucleotides, the nucleic acid template is selectively cleaved at designated positions by RNase A/T1. Mapping the resulting positional emission signatures, combined with ESI-MS analysis, allows for the discovery of cleavage hotspots. **b,** Proposed dual-path color-changing mechanism. The green-to-red conversion arises from two concurrent pathways triggered by host cleavage. Path 1 (Green-to-Dark): Dissociation of the green-emissive Ag_10_ species into non-emissive silver species (Ag_0-3_). Path 2 (Dark-to-Red): Relaxation and rearrangement of a non-emissive “dark precursor” (Ag_14_) into a highly red-emitting species (Ag_13_). **c,** Expansion of the Subak family. The original Subak (for DNase I/Cas12a) is engineered to create a family of non-FRET substrates, including rSubak-1 (RNase activity), rSubak-2 (CRISPR/Cas13a-based detection), and pcSubak (photo-cleavable linker for UV light-triggered switching). **d,** Target concentration-dependent fluorescence spectra were measured under 280 nm excitation. Fluorescence emission spectra show a distinct 95 nm red shift (from 530 nm to 625 nm) as the target concentration increases, providing a clear ratiometric readout. **e,** Key applications and performance characteristics. Subak substrates offer a cost-effective alternative to commercial substrates (e.g., RNaseAlert), maintain high sensitivity and robustness in complex biological matrices (up to 10% human serum), and facilitate point-of-care diagnostics through naked-eye colorimetric detection.

Here we investigate the cluster transformation process using site-specific cleavage of the nucleic acid templates (**Fig. 2**). By systematically substituting selected deoxyribonucleotides with ribonucleotides along the template sequence, we direct cleavage by an RNase A/T1 mixture and map sites that maximize the color changes after template cleavage. Through combined analysis of absorption and emission spectra, together with electrospray ionization mass spectrometry (ESI-MS, **Fig. 3**), we identify two color-switching hotspots (C17rC and C26rC) and discover distinct AgNC species that respond differently to template cleavage. Our data support a model in which cleavage drives loss of a green-emitting cluster (Ag_10_^6+^) and activation of a red-emitting cluster (Ag_13_^9+^) through reorganization of silver-nucleobase coordination, rather than a single, linear fragmentation pathway. This mechanism represents a previously unrecognized transformation pathway, distinct from the cluster fragmentation hypothesis in which a larger green-emitting cluster breaks down to generate a smaller red-emitting cluster^1^. Additional color conversions upon template cleavage may also exist, relying on cluster transformation mechanisms not addressed in this study. UV-initiated site-specific cleavage using a photo-cleavable (PC) linker recapitulates the same switching, further supporting a cleavage-governed mechanism with no contribution from direct enzyme-cluster interactions.

**Fig 2:**
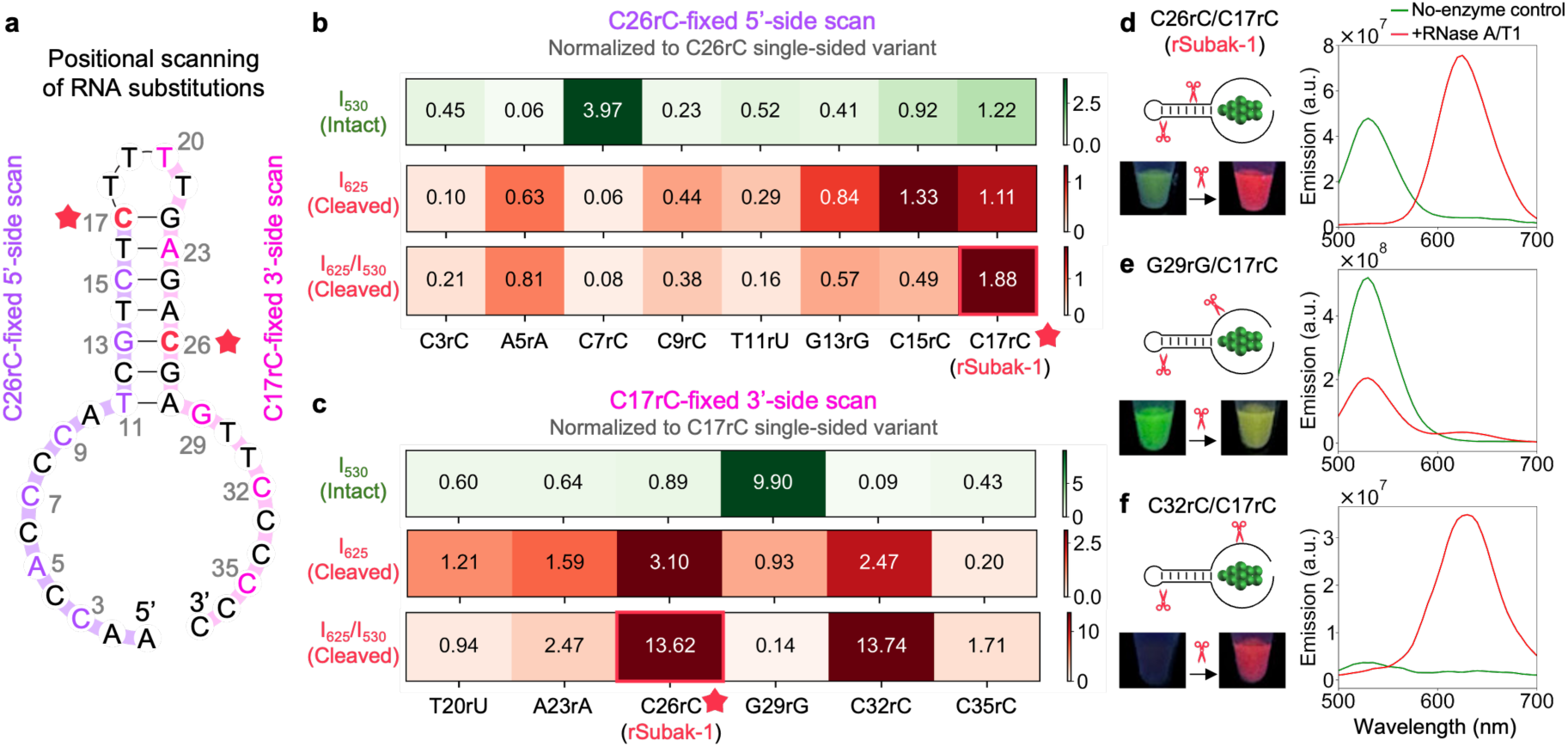
Systematic positional scanning of RNA substitutions to identify cleavage hotspots. **a,** Strategy for positional scanning of ribonucleotide substitutions. Ribonucleotides (rC, rA, rG, or rU) were systematically introduced into the Subak template at specific positions to map the sensitivity of the green-to-red color transformation to site-specific cleavage. Two scanning libraries were constructed: a 5’-side scan (purple) with C26rC fixed, and a 3’-side scan (pink) with C17rC fixed. **b, c,** Heatmaps of fluorescence responses for the 5’-side (**b**) and 3’-side (**c**) scans. The green fluorescence intensity of the intact probe (I_530_, Intact) and the red fluorescence intensity after RNase A/T1 cleavage (I_625_, Cleaved) are normalized to the single-substituted control (C26rC or C17rC). In both scans, the double substitution at C17 and C26 (designated as rSubak-1) yielded the most pronounced ratiometric increase, identifying these positions as critical cleavage hotspots. **d-f,** Representative fluorescence spectra (280 nm excitation) and tube images of selected variants. C26rC/C17rC (rSubak-1) exhibits a nearly complete transition from green (I_530_) to red (I_625_) emission upon digestion (**d**). G29rG/C17rC retains dominant green fluorescence after digestion with minimal red conversion, indicating that cleavage at these sites does not trigger the cluster transformation (**e**). C32rC/C17rC shows high red emission after digestion but suffers from poor initial green fluorescence, leading to an inefficient switch (**f**).

**Fig. 3:**
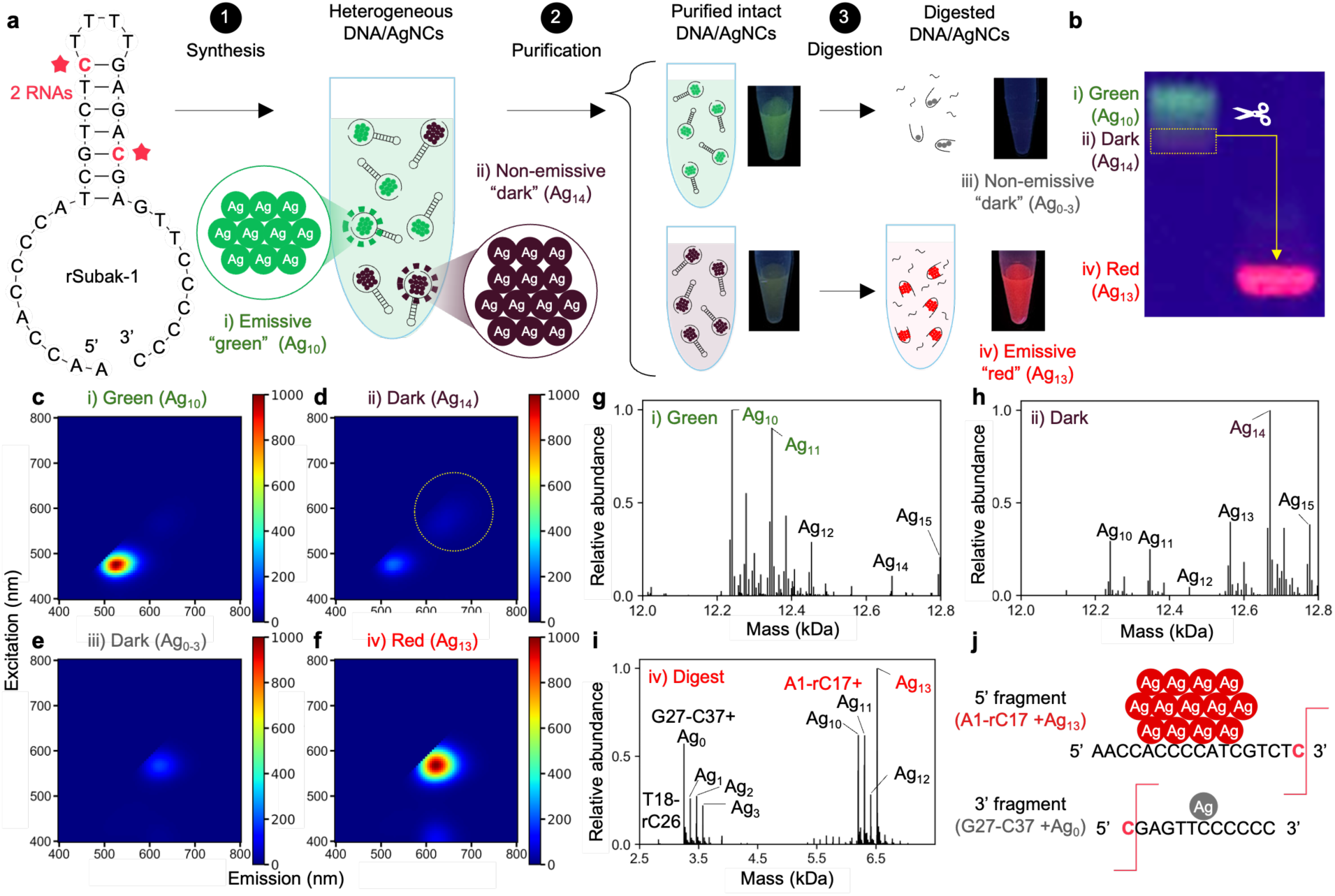
ESI-MS-based identification of cluster species and elucidation of the dual-path transformation mechanism. **a,** Experimental workflow for the synthesis, purification, and site-specific digestion of rSubak-1. Synthesis produces a heterogeneous mixture of AgNCs, which are separated into an emissive green species (Ag_10_) and a non-emissive dark precursor (Ag_14_) via purification. Digestion reveals two distinct fates: the green species dissociates into non-emissive fragments, while the dark precursor transforms into a red-emitting species (Ag_13_). **b,** Native PAGE gel image showing the physical separation of the green and dark bands. Extraction and subsequent digestion of the dark band specifically yield red fluorescence, confirming its role as the precursor for the 625 nm emission. **c-f,** Excitation-emission matrix (EEM) spectra of the isolated components before and after digestion: (i) purified green species, (ii) purified dark precursor, (iii) digested green species (non-emissive), and (iv) digested dark precursor (red-emitting). The red species formed after cleavage exhibits a characteristic peak at 625 nm. **g-i,** ESI-MS spectra providing definitive identification of the cluster compositions. The purified green species is dominated by Ag_10_ and Ag_11_ clusters (**g**), whereas the non-emissive dark precursor consists primarily of Ag_14_ clusters (**h**). Upon digestion (**i**), the Ag_14_ species form the red-emitting Ag_13_ cluster. **j,** Schematic mapping of cluster-fragment products, identifying the 5’ fragment (A1-rC17) as the host for the red-emitting Ag_13_ cluster and the 3’ fragment (G27-C37) as associated with non-emissive silver atoms.

Leveraging the insights gained from the site-specific cleavage results, we further develop two new Subak substrates, termed rSubak-1 and rSubak-2, for RNase A and Cas13 activity sensing (**Fig. 4**). In amplification-free CRISPR/Cas13a assays, rSubak-2 achieves limits of detection (LoD) of 0.7 pM and 0.3 pM for SARS-CoV-2 and influenza A virus (A/H5N1), respectively, representing >300-fold improved sensitivity relative to the commercial FRET substrate, RNaseAlert (LoD 130-250 pM), at substantially lower substrate cost (one seventy-sixth). Owing to its ratiometric readouts, rSubak-2 provides a broad quantifiable dynamic range (1 pM to 100 nM) and enables accurate quantification of viral loads below 250 pM that are undetectable by RNaseAlert. Moreover, rSubaks perform robustly in complex biological matrices (e.g., 10% human serum) and retain activity after lyophilization and rehydration (**Fig. 5**), supporting their use in the field^22^.

**Fig. 4:**
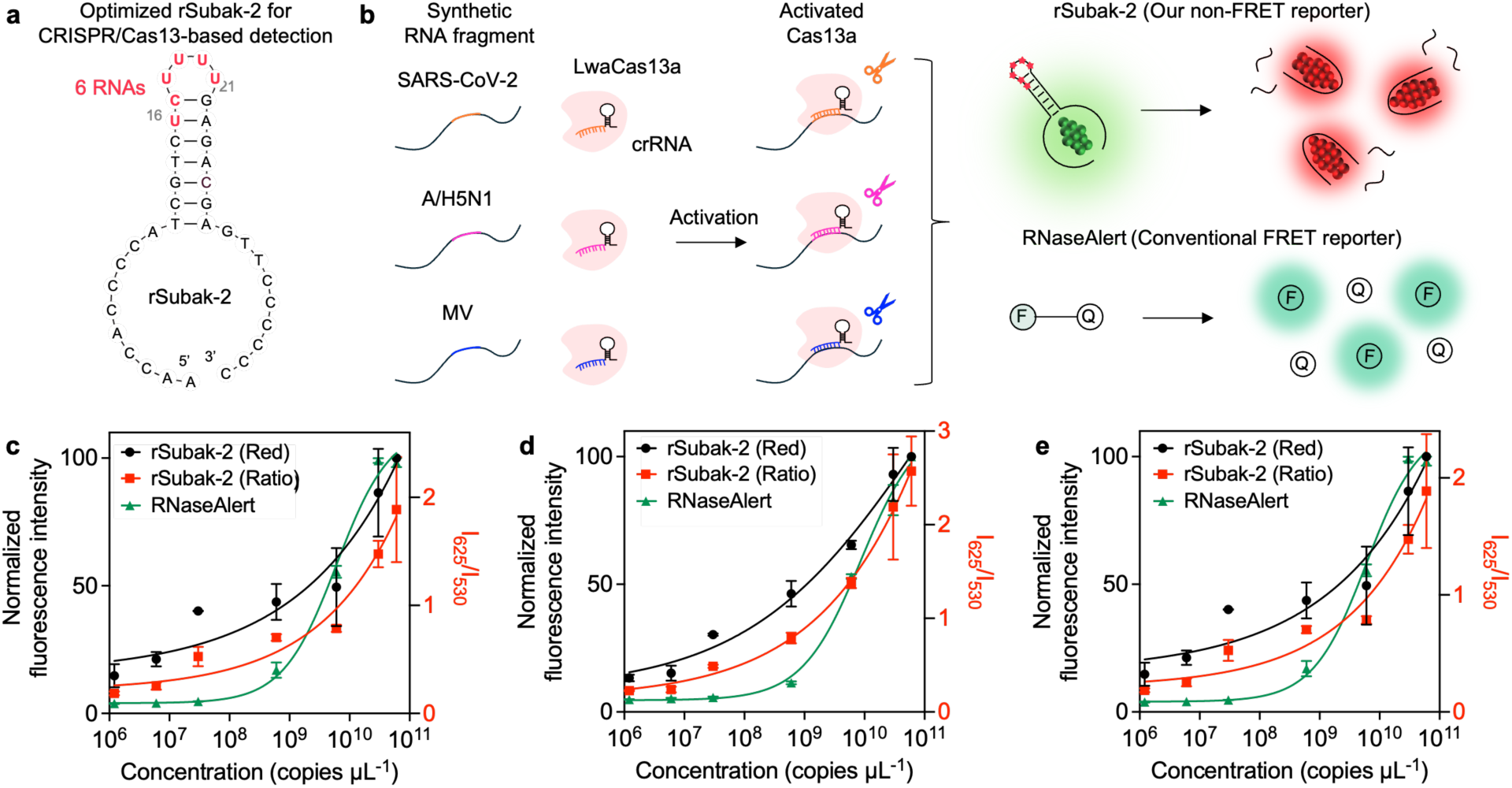
CRISPR/Cas13a-based quantitative detection of viral RNAs using the optimized rSubak-2 substrate. **a,** Secondary structure of the optimized rSubak-2 substrate. The poly-uridine (poly-U) sequence (rU16-rU21) in the loop region, and the cytosine at position 17 is substituted with a ribonucleotide (rC17) to facilitate the trans-cleavage of activated LwaCas13a. **b,** Schematic illustration of the CRISPR/Cas13a detection assay. Target viral RNA fragments (SARS-CoV-2, A/H5N1, and MV) activate the LwaCas13a-crRNA complex, triggering the *trans-cleavage* of substrates. Note that only the emissive species are depicted in the schematic. **c-e,** Normalized target-dependent fluorescence responses for SARS-CoV-2 (**c**), A/H5N1 (**d**), and MV (**e**) synthetic RNA fragments. The fluorescence intensity at 625 nm (rSubak-2, black circles) and 520 nm (RNaseAlert, green triangles) are normalized to 100 at the maximum concentration for each target. Data are presented as the average of n=2 independent replicates. Emission spectra were acquired under 280 nm excitation.

**Fig. 5:**
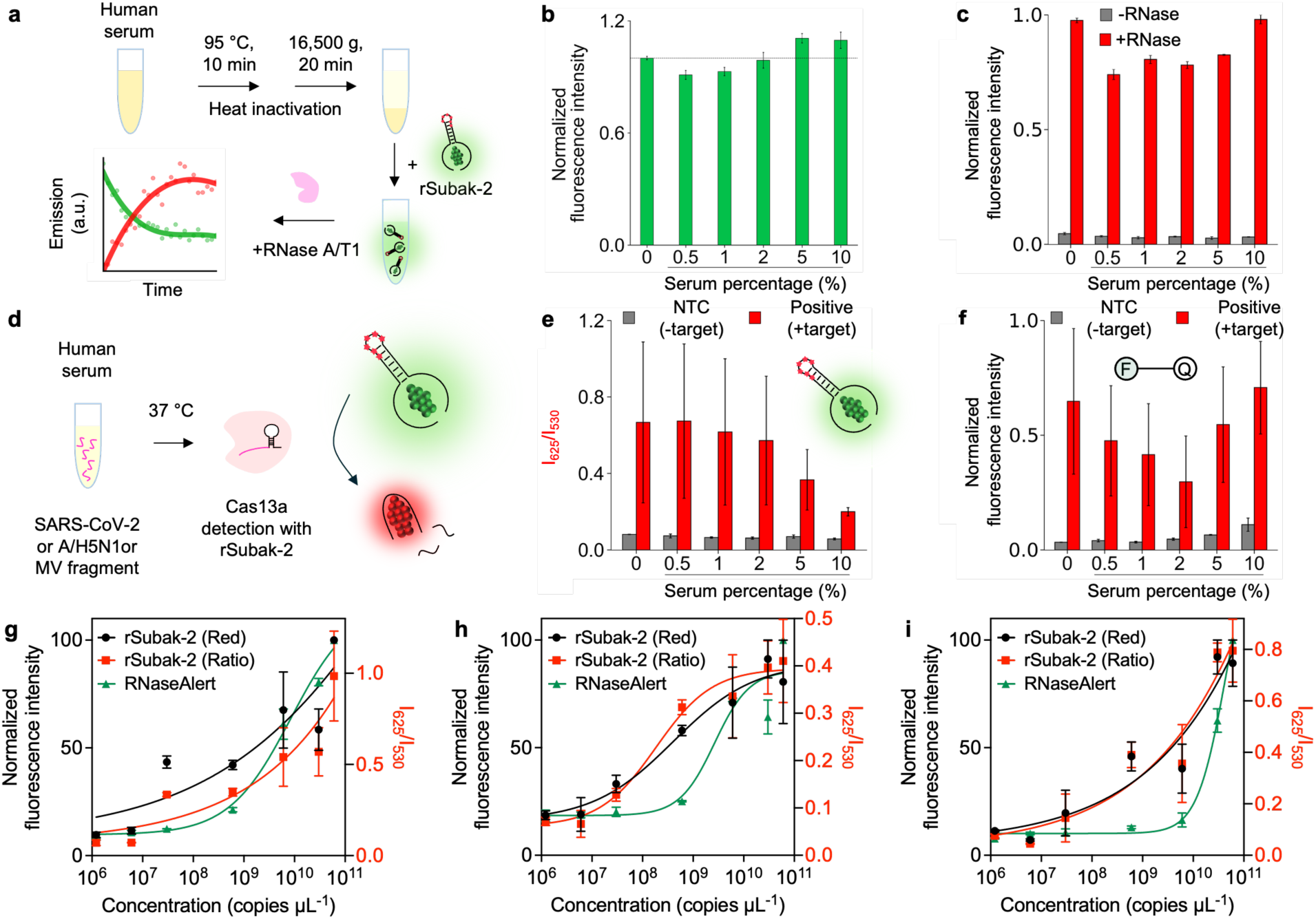
Evaluation of rSubak-2 performance and stability in complex biological matrices. **a,** Schematic of the experimental setup for testing rSubak-2 stability and activity in human serum. Serum samples were heat-inactivated (95°C, 10 min) and centrifuged before being spiked with the substrate and target. **b, c,** Stability and reactivity of rSubak-2 across varying serum percentages (0-10%). The normalized green fluorescence (I_530_) remains stable in the absence of RNase (**b**), while addition of RNase A/T1 triggers a robust red fluorescence response (I_625_) across all tested serum concentrations (**c**). **d,** Schematic of the CRISPR/Cas13a-based viral RNA detection assay conducted in human serum. **e, f,** Comparison of ratiometric rSubak-2 response (I_625_/I_530_, **e**) and conventional FRET-based RNaseAlert response (**f**) in serum-spiked samples. rSubak-2 maintains a clearer signal-to-background separation compared to the FRET substrate in complex environments. **g-i,** Normalized target-dependent fluorescence responses for SARS-CoV-2 (**g**), A/H5N1 (**h**), and MV (**i**) synthetic RNA fragments in 2% human serum. The fluorescence intensity at 625 nm (rSubak-2, black circles) and 520 nm (RNaseAlert, green triangles) are normalized to 100 at the maximum concentration for each target. Data are presented as the average of n=2 independent replicates. Emission spectra were acquired under 280 nm excitation. Note that only the emissive species are depicted in the schematics for clarity.

By incorporating ribonucleotides into Subak templates, we not only expand the Subak family into substrates for RNase activity sensing but also gain mechanistic insights into cleavage-governed AgNC transformation (**Fig. 6**), paving the way for rationally designing novel substrates that can effectively detect the activities of other cleavage enzymes, such as proteases and cellulase. Moreover, our non-FRET substrates are prepared in a one-pot reaction, where chromatographic purification is not required for routine assay use. Collectively, these results highlight the robustness and versatility of Subak-family substrates as low-cost, complex-matrix-tolerant alternatives to conventional FRET probes for bioassays and diagnostics.

**Fig. 6:**
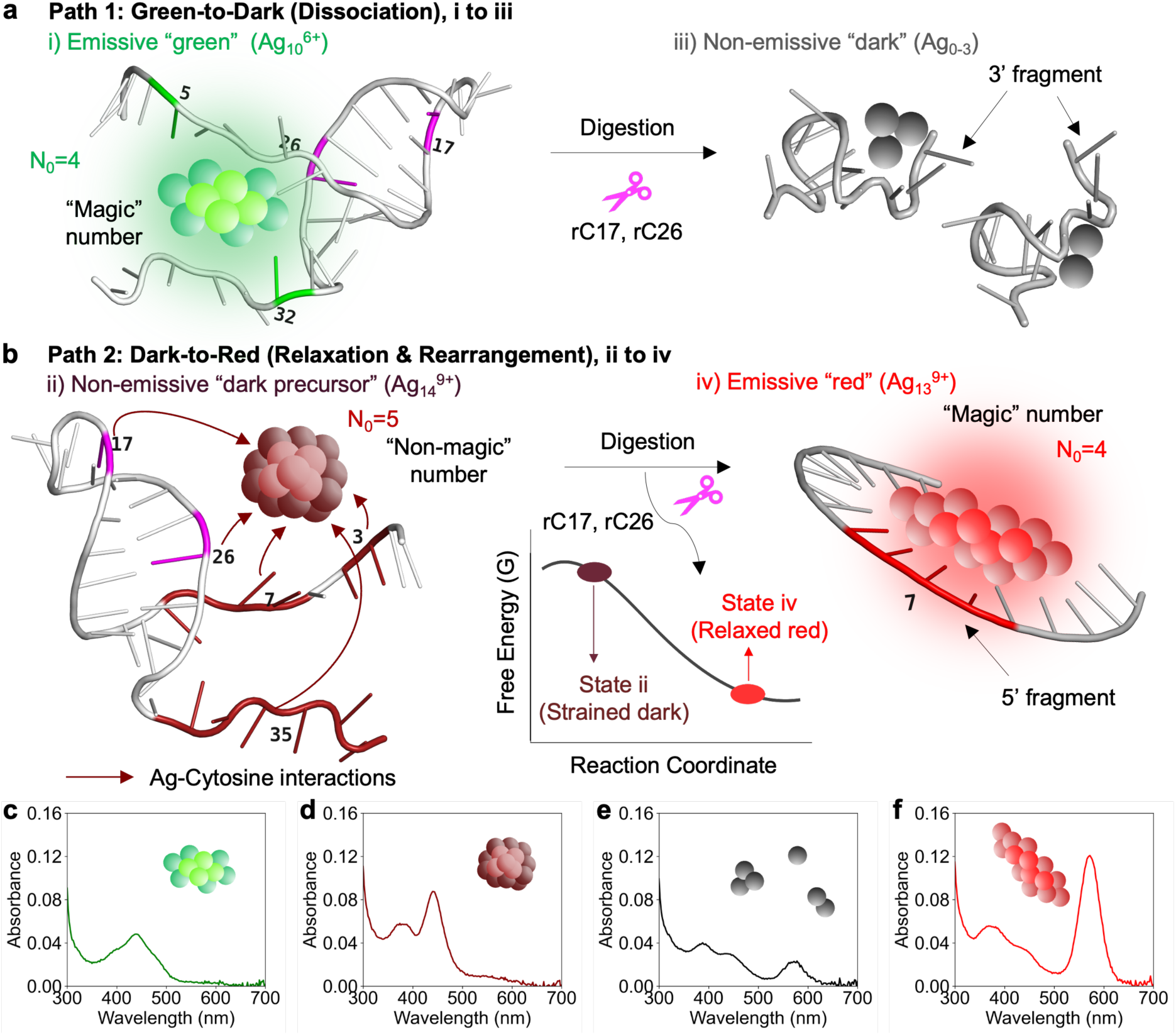
Proposed dual-path mechanism reinforced by absorbance characteristics. **a, b,** Structural models and schematic pathways illustrating the green-to-dark dissociation (**a**) and dark-to-red relaxation (**b**) mechanisms. Upon site-specific cleavage at rC17 and rC26, the emissive green Ag_10_ species (Path 1) dissociates into non-emissive Ag_0-3_ fragments. Simultaneously, the non-emissive dark precursor (Ag_14_, Path 2) undergoes structural relaxation and rearrangement into a thermodynamically stable, red-emitting Ag_13_ cluster. **c-f,** Absorbance spectra providing final spectroscopic validation for the four corresponding states: (i) pre-cleavage green Ag_10_ (**c**), (ii) pre-cleavage dark Ag_14_ (**d**), (iii) post-cleavage dark species (**e**), and (iv) post-cleavage red species (Ag_13_, **f**). The emergence of a distinct absorption peak at 572 nm in (**f**) serves as a complementary indicator for the formation of the red-emitting cluster, consistent with the proposed transformation model.

## RESULTS

### Subak as a versatile non-FRET platform for cleavage-based detection

**Fig. 1** provides an overview of this study. To gain mechanistic insights into the cleavage-induced color transformation of Subak, here we systematically substituted selected deoxyribonucleotides with ribonucleotides along the Subak-2 template sequence, which was a DNase substrate described in our previous report^1^ (**Fig. 1a** and **Supplementary Table 1**). These engineered RNA sites enabled site-specific cleavage by an RNase A/T1 mixture (except at rA). The resulting DNA-RNA chimeric constructs addressed a key limitation in studying the DNase substrate Subak, namely the lack of control over DNase cleavage sites. By mapping color transformations to defined cleavage positions and integrating these results with ESI-MS analysis, mechanistic insights into the structural evolution of AgNCs were obtained (detailed below). Based on these findings, we propose a dual-path color-changing mechanism (**Fig. 1b**). Cleavage at specific positions, termed hotspots, led to effective destabilization of the green-emitting Ag ^6+^ cluster (Path 1), while the weakly emissive precursor underwent structural rearrangement to generate a strongly emissive red Ag_13_^9+^ cluster (Path 2). Guided by the site-specific cleavage results, we designed two RNA-incorporated Subak substrates (rSubaks) to detect the activities of distinct RNases: rSubak-1 for RNase A and rSubak-2 for activated Cas13a (**Fig. 1c**, **Supplementary Table 1** and **Supplementary Fig. 1**). In particular, rSubak-1 responded strongly to the RNase A/T1 mixture, exhibiting a clear green-to-red emission shift upon cleavage (530 to 625 nm, **Fig. 1d**). Additionally, a photo-cleavable-linker-incorporated Subak (pcSubak) was also constructed. Upon photolysis, pcSubak likewise exhibited a green-to-red color conversion, further supporting the cleavage-driven transformation mechanism (**Supplementary Fig. 2**). Compared with the commercial FRET substrate RNaseAlert, our Subak platform offers several advantages, including substantially lower cost ($1 vs $76 per nanomole), a one-pot reaction workflow, much reduced no-target-control background signals in 10% serum and a readily visible naked-eye color change which enables quantitative ratiometric sensing (**Fig. 1e, Supplementary Table 2,** and **Supplementary Fig. 3**).

### Mapping and structural profiling of DNA-RNA chimeras under site-specific cleavage

Because the initial single-site cleavage investigation produced only moderate color transformation, likely due to incomplete dissociation of cleavage fragments (**Supplementary Fig. 4-6**), here we focused on screening double-cleavage sites. We selected two preferred cleavage positions (17 and 26) identified in the previous DNase I experiments^1^ as starting points (**Fig. 2a**). Prior ESI-MS analysis indicated that the AgNC remained associated with the 5′ fragment after cleavage. Accordingly, we fixed one cleavage site at position 26 (C26rC, denoting substitution of deoxycytidine at position 26 with cytidine) and systematically introduced a second cleavage site along the 5′ region, generating 8 double-cleavage chimeric constructs (**Fig. 2b**). Similarly, by fixing the cleavage site at C17rC and scanning along the 3′ region, 6 additional double-cleavage constructs were produced (**Fig. 2c**). As expected, all 14 constructs exhibited varying degrees of green-to-red color transformation upon treatment with the RNase A/T1 mixture. However, substantial differences were observed among them. Because the RNase A/T1 mixture hydrolyzes RNA specifically at the 3′ end of rU, rC, and rG residue^23^, two chimeric constructs effectively underwent only single-site cleavage under these conditions (A5rA/C26rC and A23rA/C17rC).

Using the single-cleavage constructs C26rC and C17rC as benchmarks, we compared fluorescence outputs. Some double-cleavage constructs exhibited stronger initial green emission (I_525_, top row in **Fig. 2b**), but their post-cleavage red emission (I_630_, middle row) was markedly weaker. For instance, C7rC/C26rC and G29rG/C17rC samples displayed normalized (I_525_, I_630_) values of (3.97, 0.06) and (9.90, 0.93), respectively, reflecting poor AgNC transformation efficiency. In contrast, C32rC/ C17rC exhibited the strongest red emission after cleavage, yet its initial green emission was very low (**Fig. 2c**), with normalized (I_525_, I_630_) values of (0.09, 2.47), indicating strong environmental sensitivity of AgNC fluorescence. When evaluating these double-cleavage constructs using two criteria: (1) a large ratiometric change (I_525_/I_630_; bottom rows in **Fig. 2b,c**) and (2) a visually distinct color conversion before and after cleavage (**Fig. 2d-f**), C17rC/C26rC emerged as the best-performing construct (**Supplementary Fig. 7**).

Given that cytosine and guanine exhibit the strongest interactions with silver^24–29^, it is unsurprising that hotspots showing the largest fluorescence or color changes correspond to C and G positions (e.g., C17rC, C26rC, G29rG and C32rC). Interestingly, G29rG substitution markedly enhanced pre-cleavage green emission (**Fig. 2e**), whereas C32rC nearly abolished it (**Fig. 2f**). These results suggest that AgNC fluorescence is sensitive not only to nucleobase identity but also to sugar composition, potentially through altered AgNC binding sites within the template^13,30^ and changes in dipolar interactions^25^. Close examination of the three representative constructs (**Fig. 2d-f**) led us to hypothesize that the C32rC/C17rC sample contains a low-emissive AgNC species that can be efficiently converted into a red-emitting cluster upon template cleavage. In contrast, the G29rG/C17rC template appears to preferentially support synthesis of the green-emitting cluster but not the weakly emissive precursor. This hypothesis is subsequently confirmed by gel electrophoresis (see below).

Importantly, the double-cleavage experiments not only mapped color transformations to defined cleavage sites but also clarified why double-cleavage construct C17rC/C26rC represents the optimal design: it balances the synthesis yields of the green-emitting AgNC species and the nearly non-emissive precursor. The coexistence of these two species underlies the pronounced color transformation observed upon double-cleavage of the C17rC/C26rC construct.

Substitutions within the stem region (positions T11-C17 and G22-A28) generally reduced green fluorescence prior to cleavage (e.g., G13rG/C26rC and A23rA/C17rC). This effect likely arose because RNA-DNA base pairing is stronger than DNA-DNA pairing, thereby further stabilizing the stem. A moderately stable stem in the hairpin might therefore favor a higher synthesis yield of the green-emitting species. In contrast, substitutions within the 5′ binding arm (positions A1-A10) abolished red emission after cleavage (e.g., C3rC/C26rC and C7rC/C26rC; **Fig. 2b-c**), indicating that the integrity of the A1-C9 segment is critical for formation and stabilization of the red-emitting AgNC following cleavage. Notably, the strong red emission observed in the A5rA/C26rC construct arose because the RNase A/T1 mixture cleaves at C26rC but not at A5rA, thereby preserving the A1-C9 segment.

Because G29rG substitution significantly enhanced pre-cleavage green emission while maintaining strong post-cleavage red emission, we hypothesized that a triple-substitution construct (C17rC/C26rC/G29rG) might further amplify the green-to-red transformation. Indeed, this triple-cleavage construct increased the initial green signal in the intact state. However, its ratiometric performance was inferior to that of C17rC/C26rC (I_625_/I_530_ = 12.7 vs. 48.3), due to substantial residual green fluorescence after cleavage (**Supplementary Fig. 8**).

The top-performing construct from the site-specific cleavage screening, C17rC/C26rC, was subsequently renamed as rSubak-1 in the following experiments. This optimized construct exhibited ∼2.5-fold higher red emission after cleavage compared with the single-cleavage control C26rC (**Supplementary Fig. 7**). As a DNA-RNA chimeric construct, rSubak-1 remained susceptible to DNase digestion (**Supplementary Fig. 9**). However, RNase A/T1-mediated cleavage produced ∼6.2-fold stronger red emission than DNase I cleavage, highlighting the advantage of site-specific cleavage in maximizing color transformation efficiency (**Supplementary Fig. 9**).

### Mass spectrometry characterization of cluster stoichiometry

To elucidate the structural basis of the fluorescence transition, we performed electrospray ionization mass spectrometry (ESI-MS) on gel-purified rSubak-1 before and after RNase A/T1 cleavage (**Fig. 3**). Intact rSubak-1 displayed two distinct substates: a green-emitting species and a nearly non-emissive “dark” species. Upon RNase A/T1 cleavage, these substates underwent distinct transformation: the green substate gradually lost its fluorescence, while the dark substate converted into a red-emitting species (**Fig. 3a,b**).

We further analyzed the cleavage behaviors and optical properties using gel electrophoresis coupled with excitation-emission matrix (EEM) and ESI-MS (**Fig. 3c-i**). Before cleavage, rSubak-1 showed two bands corresponding to the green and dark substates, which exhibited distinct fluorescence signatures (**Fig. 3c,d**). Analysis of the excised gel bands by ESI-MS revealed distinct Ag_n_ distributions associated with each population: the green band was enriched in smaller Ag clusters (predominantly N = 10-11), whereas the dark band was enriched in larger Ag clusters (N = 14) (**Fig. 3g,h**). Following cleavage, the green band’s signal disappeared (Fig. 3e), while a single, intense red-emitting band emerged (**Fig. 3f**), indicating efficient cleavage and AgNC transformation.

ESI-MS analysis clarified the stoichiometric basis of this transformation. Unlike the DNA-only system where large green-emitting Ag_13_/Ag_14_ clusters fragment into smaller Ag_10_/Ag_11_ clusters^8^, rSubak-1 operates through a distinct mechanism. While the initial green fluorescence arises from Ag_10_/Ag_11_ clusters, the final red fluorescence corresponds to a Ag_13_ cluster (**Fig. 3g-i**). Notably, this red-emitting Ag_13_ species is generated from the abundant non-emissive dark precursor (Ag_14_) via a minimal stoichiometric shift (Ag_14_ to Ag_13_), rather than from fragmentation or consolidation of the green clusters. Schematic mapping confirmed that the Ag_13_ cluster is hosted in the 5’ fragment (A1-C17rC), while the 3’ fragment carries only minimal silver atoms and lacks the coordination sites required for an emissive cluster (**Fig. 3j**). Analysis of the isotopic distributions reveals the charge states of green-emitting, nearly non-emissive, and red-emitting clusters to be Ag_10_^6+^, Ag_14_^9+^, and Ag_13_^9+^, respectively (**Supplementary Fig. 10-12**).

These findings provide direct evidence that fluorescence switching in rSubak is not driven by extensive silver loss or cluster fragmentation. Instead, the cleavage of the DNA/RNA stem triggers a transformation in the coordination environment of the dark precursor Ag_14_^9+^. While a minimal stoichiometric change (Ag_14_^9+^ to Ag_13_^9+^) is observed, the backbone cleavage likely reconfigures the silver-nucleobase interaction footprint, enabling the transformation from a non-emissive state to a red-emitting state. A detailed structural model explaining this geometry-driven transition, distinct from the fragmentation model, is proposed in the Discussion section.

### Engineering rSubaks for efficient CRISPR/Cas13a recognition

We next adapted the system for CRISPR/Cas13a-based RNA detection assays, where an RNA-guided RNase Cas13a enzyme collaterally cleaved surrounding RNA substrates upon target recognition (i.e., *trans*-cleavage)^31–33^. We first evaluated rSubak-1, which contains two ribonucleotide substitutions (C17rC/C26rC, **Fig. 3a**). Although rSubak-1 responded efficiently to RNase A/T1 cleavage (exhibiting a >6.2-fold increase in red emission), the same construct yielded a negligible response to Cas13a *trans*-cleavage (with no discernible signal-on response above the no-target-control, NTC, baseline; **Supplementary Fig. 13**). This poor response could stem from biochemical and structural differences among nucleases. In fact, Cas13a is substantially larger (∼144 kDa) than DNase I (39 kDa) and RNase A (11 kDa), corresponding to ∼1.5-fold and ∼2.3-fold differences in hydrodynamic diameters, respectively. As a result, the Cas13a-crRNA ribonucleoprotein (RNP) complex imposes far more stringent steric and conformational requirements on its substrate, particularly around its catalytic HEPN (higher eukaryotes and prokaryotes nucleotide-binding) nuclease domains that specifically cleave RNA linkages^34^. In contrast, RNase A can have more efficient access to mixed DNA/RNA backbones, consistent with its higher cleavage efficiency on rSubak-1. Hence, optimizing rSubak for the Cas13a system requires balancing the number of ribonucleotide substitutions for better recognition and easy cleavage, with the coordination of deoxyribonucleotides that stabilize desired AgNC species before and after cleavage.

To systematically probe this trade-off and create an effective rSubak for Cas13 *trans*-cleavage sensing, we screened a library of chimeric DNA/RNA substrates derived from the Subak-2 scaffold, which previously exhibited superior color-switching performance with Cas12a^1^. The library included variants with 2-4 consecutive ribouridines (rU2-rU4), alternating rG/rU bulges within the central stem, and 5-6 consecutive ribonucleotides without bulges (**Supplementary Table 3**). While introducing bulges could improve the local accessibility to Cas13a, these variants exhibited significantly higher pre-cleavage red emission (**Supplementary Fig. 14**). This high background was attributed to their severely compromised thermodynamic stability of the hairpin structures. The 2-bulge variant showed a ΔG_folding_ of only −3.6 kcal mol^−1^ and a melting temperature (T_m_) of 47.3 °C, compared to the highly stable intact stem (ΔG ≈ −8.2 kcal mol^−1^, T_m_ ≈ 70.1 °C). Such low stability likely induces spontaneous structural breathing near the physiological temperature (37 °C), perturbing the coordination environment to stabilize desired clusters before and after cleavage.

In contrast, variants with 5-6 consecutive ribonucleotides maintained the integrity of the stem (ΔG ≈ −8.2 kcal mol^−1^), providing the best trade-off between biochemical reactivity and photophysical performance. During optimization, we also explored a dual-sided cleavage design to further enhance post-cleavage red emission; however, while the red signal increased, it often resulted in significantly quenched pre-cleavage green fluorescence or an elevated red background fluorescence (**Supplementary Fig. 15**). In particular, the construct with six consecutive ribonucleotides from position 16 to 21 showed minimal pre-cleavage red background fluorescence and retained bright, distinct red emission after cleavage with negligible residual green signal (**Supplementary Fig. 15**). These results confirmed that expanding the RNA segment to 6-nt, without introducing destabilizing bulges, maximizes Cas13a accessibility while preserving the AgNC-stabilizing scaffold. We named the optimized construct rSubak-2 (**Fig. 4a**) and integrated it into CRISPR/Cas13a assays for amplification-free RNA viral detection.

To demonstrate the versatility of the platform, we evaluated rSubak-2 in amplification-free Cas13a assays targeting three representative RNA viral strands: SARS-CoV-2, avian influenza A H5N1 (A/H5N1), and measles virus (MV). Unlike RNaseAlert that provides only an intensity-based turn-on signal, rSubak-2 showed a clear green-to-red fluorescence transition (530 to 625 nm) upon Cas13a *trans*-cleavage, enabling ratiometric quantification (**Fig. 4b**). Across all targets, rSubak-2 showed superior sensitivity and a broader quantifiable dynamic range compared to the commercial benchmark (**Fig. 4c-e**). For SARS-CoV-2, the amplification-free LoD was as low as 0.7 pM (≈ 6.6 × 10⁵ copies µl^−1^) using the ratiometric readout (I_625_/I_530_), which was 3.7-fold more sensitive than the LoD (2.6 pM) from the single-channel readout (I_625_) and significantly lower than that of RNaseAlert (130 pM). Similarly, using the ratiometric readout rSubak-2 achieved improved LoD for A/H5N1 (0.3 pM) and MV (57 pM) sensing compared to the FRET substrate (248 pM and 152 pM, respectively) (**Fig. 4d,e**). Beyond sensitivity, rSubak-2 yielded a 19-fold ratiometric fluorescence fold change upon cleavage by Cas13a, a 1.3-fold improvement over the original Subak-2 previously reported for Cas12a-mediated detection^1^. Furthermore, by expanding the quantifiable window across 5 orders of magnitude (from 1 pM to 100 nM), rSubak-2 enabled precise quantification of viral loads that remained undetectable by RNaseAlert, which was insensitive when the target strand concentration was below 300 pM. Notably, these performance gains came at a fraction of the cost. rSubak-2 could be synthesized for ∼$1 per nanomole, as compared to ∼$76 per nanomole of RNaseAlert when purchased from the vendor, supporting scalable deployment for mass testing at resource-limited settings^35^.

### Comparing performance of rSubak with RNaseAlert in diluted serum

We quantitatively assessed the robustness of rSubak-2 in complex biological matrices using heat-inactivated human serum (**Fig. 5a**). Following the protocol outlined in literature^34,35^, serum samples were pre-treated (95 °C, 10 minutes) to inactivate intrinsic nucleases. To validate substrate stability in the absence of target RNA, background fluorescence was monitored across all tested serum dilutions (0-10% v/v). After a 30-minute incubation, the green emission intensities (I_530_) remained stable, maintaining a range of 0.75-1.11 relative to the buffer baseline (normalized to 1.0 at 0% serum, **Fig 5b**). Although a slight signal enhancement (up to ∼10%) was observed in 5% serum, likely due to cation-mediated stabilization of the DNA/AgNCs scaffold^36–38^, the overall background signal remained within 11% of the buffer baseline (0%, **Fig. 5b**), confirming minimal non-specific activation.

To confirm that the green-to-red color switching remains functional in serum, we tested rSubak-2 with a positive control (RNase A/T1 mixture), a robust cleavage enzyme. The post-cleavage red emission (I_625_) was consistently observed across all tested serum concentrations (0-10%), maintaining over 75% of the buffer-only intensity (**Fig. 5c**). This result demonstrates that serum components do not interfere with the cleavage-induced transformation of the AgNC, ensuring the feasibility of rSubak-2 substrates in clinical matrices.

We then examined Cas13a-mediated detection directly in serum (**Fig. 5d**). As serum concentration increased from 0% to 10%, the ratiometric signal (I_625_/I_530_) decreased from ∼0.67 to ∼0.20. This reduction likely results from the combined effects of enhanced green fluorescence of the intact AgNCs and partial inhibition of Cas13a activity by serum proteins (**Fig. 5e**). Nevertheless, the system maintained a clear distinction between positive and NTC samples; even in 10% serum, the positive signal was 3.5-fold higher than the NTC baseline, enabling reliable target discrimination. In contrast, RNaseAlert failed at serum levels above 5% due to non-specific activation by endogenous nucleases^39^ (**Fig. 5f**). This superior stability stems from the structural design of the substrates: while RNaseAlert is an all-RNA construct highly susceptible to ubiquitous serum RNases, rSubak-2 is a DNA-RNA chimeric construct. By embedding the RNA cleavage site within a DNA-stabilized scaffold, rSubak-2 effectively shields the substrates from non-specific degradation while remaining highly responsive only to the target-activated Cas13a complex. To ensure diagnostic integrity and consistency, 2% serum was selected as the optimal condition^33^ in the rest of the evaluation.

In testing spike-in samples within 2% serum, rSubak-2 successfully detected synthetic SARS-CoV-2, A/H5N1, and MV viral RNA strands with a strong concentration-dependent response (**Fig. 5g-i**). Across all targets, the ratiometric LoD were 13 pM (SARS-CoV-2, **Fig. 5g**), 0.4 pM (A/H5N1, **Fig. 5h**), and 16 pM (MV, **Fig. 5i**). These values are consistently lower than those of the RNaseAlert (31 pM, 253 pM, and 1.0 nM, respectively), demonstrating the superior performance of rSubak-2 in complex serum environments. Finally, to evaluate field deployability, reagents were lyophilized and stored at room temperature (**Supplementary Fig. 16**). Upon rehydration, the assay preserved fluorescence kinetics and end-point intensity, confirming that rSubak is a robust, ready-to-use platform for point-of-care diagnostics.

## Discussion

In this study, we introduce rSubak, a versatile and cost-effective RNase substrate that exhibits pronounced green-to-red color conversion upon nucleic acid template cleavage. Our results indicate that incorporation of two or more ribonucleotide cleavage sites is required to achieve maximal color switching and cleavage efficiency especially when a bulky RNase such as Cas13a is used. Multiple cleavage events likely promote complete dissociation of the 3′ arm from the 5′ arm (where AgNC resides after cleavage), thereby facilitating reorganization of the nucleobase environment surrounding the AgNC and ultimately leading to strong red fluorescence emission. Notably, photolytic cleavage also produces a red-emissive state, confirming that the color-switching behavior is governed by cleavage-induced local structural rearrangements of the scaffold rather than direct enzyme-cluster interactions.

Our findings offer a refined perspective on the underlying mechanism of this transition, moving beyond the traditional cluster fragmentation model. By identifying two specific cleavage hotspots (C17rC, C26rC), we characterized a dual-path transition involving two coexisting cluster populations that respond differently to host cleavage (**Fig. 6a,b**). The first pathway (Path 1) involves the green-emitting Ag_10_^6+^ species. In the intact scaffold, this cluster adopts a geometry that supports strong green emission peaked at 530 nm, which is also commonly observed by other DNA/AgNC researchers^28,40–42^. However, upon site-specific cleavage of the DNA/RNA stem, the coordination environment is disrupted, leading to cluster fragmentation or dissociation, which effectively “turns off” the green fluorescence.

The second and more critical pathway (Path 2) explains the emergence of red fluorescence from the dark Ag_14_^9+^ precursor. A key question lies in why the Ag_14_ species remains non-emissive (dark) in the intact state despite its high silver nuclearity. We hypothesize that the Ag_14_ cluster is kinetically trapped within the rigid, cytosine-rich hairpin. In this constrained environment, the cluster is forced into a compact, spherical-like geometry, as evidenced by the multiple peaks at shorter wavelengths (440 nm) in its absorption spectrum (**Fig. 6c**), which resemble the absorption peaks observed in other AgNCs that are known to carry a spherical shape^43^.

The “switching” event occurs when cleavage at the C17rC and C26rC hotspots relieves this structural strain. This scaffold relaxation allows the silver core to undergo a structural rearrangement into a thermodynamically favored, rod-like Ag_13_^9+^ geometry. Remarkably, ESI-MS confirms that this transition involves a minimal stoichiometric shift (loss of only one silver atom, from Ag_14_ to Ag_13_), which proves that the red emission originates from a geometry-driven reconfiguration rather than extensive fragmentation (**Fig. 3i, 6f**). This transition to a rod-like shape is confirmed by the emergence of a dominant, single absorption peak at 572 nm, which resembles the single-peak signature observed in other AgNCs that are known to be rod shaped^43^. This electronic activation aligns with the scaling law of cluster size (E_gap_ μ N^−1/3^), explaining why the larger Ag_13_ core exhibits a significantly redshifted emission (625 nm) relative to the smaller Ag_10_ species (530 nm)^42,44,45^. This scaffold-relaxation hypothesis also interprets the faster migration of the Ag_14_ precursor in the native PAGE (**Fig. 3b**), a result consistent with a hyper-compact, globular folding of the DNA host, which effectively reduces its hydrodynamic radius compared to the green species^12,24^. Such evidence underscores that ratiometric switching involves a significant geometric transition driven by the structural reorganization of this scaffold.

Furthermore, this study extends our understanding of RNA-AgNC interactions. Our results indicate that ribonucleotide substitutions, even at minimal levels, can subtly redefine the nucleobase-silver interaction footprint. This highlights that the additional 2’-hydroxyl groups and the distinct conformational flexibility of RNA can be strategically utilized to modulate AgNC coordination pathways. Mechanistically, this insight was instrumental in optimizing the substrate for CRISPR/Cas13a-based detection. Given the larger size of Cas13a (∼144 kDa) compared to RNase (11 kDa), we found that incorporating consecutive ribonucleotides into the stem was essential to create an RNA-accessible region while preserving the DNA rigidity required for AgNC stabilization. Although the optimized rSubak-2 balances these factors effectively and confines the color-switching event to a single, well-defined nucleation site, there remains room for further improvement.

Functionally, rSubak-2 demonstrated amplification-free RNA viral detection performance that is competitive with commercial FRET-based substrates, yet at a significantly lower synthesis cost. The assay achieved picomolar sensitivity for targets such as SARS-CoV-2 and A/H5N1, maintained robust performance in complex matrices including human serum, and remained functional after lyophilization. Building upon these results, future iterations could couple rSubak with upstream isothermal amplification methods (e.g., RPA^46,47^ or LAMP^48,49^) or integrate it with smartphone-based readout devices^46,50^ utilizing its ratiometric output. Broadly, the mechanistic insights gained here suggest a general guideline for engineering DNA/AgNCs substrates. Ultimately, by elucidating the structural interplay between chimeric scaffolds and silver clusters, this work establishes a rational design strategy for developing programmable, geometry-driven fluorescent substrates that could be extended other polymer digestion enzymes, including proteases and cellulases, as AgNCs have also been prepared in cellulose^51^ and peptide^52^.

## MATERIALS AND METHODS

### DNA/AgNCs preparation

Subak substrates were prepared in hairpin DNA templates using the conventional nucleation-and-reduction process to form AgNCs in DNA. The oligonucleotides (Integrated DNA Technologies, IDT) were suspended at 500 µM in DNase-free water and then thawed in water at 70 °C for 2 min and cooled for at least 5 min. To make 1 ml of solution, 80 µl of 500 µM DNA was added to 100 µl of 200 mM sodium phosphate buffer (SPB, pH 7.4) with 748 µl of nuclease-free water. The solution was then mixed with 48 µl of 10 mM silver nitrate (AgNO_3_, cat. no. 204390, Sigma-Aldrich) solution, and then vortexed and centrifuged for 1 min at 14,000g. After equilibration for 15 min, the mixture was reduced with 24 µl of 10 mM freshly prepared sodium borohydride (NaBH_4_, cat. no. 480886, Sigma-Aldrich) to form the DNA/AgNCs. The solution was then vortexed and centrifuged again for 1 min at 14,000 g. The final concentrations were 40 µM DNA, 20 mM SPB (pH 7.4), 480 µM AgNO_3_ and 240 µM NaBH_4_. The product was stored at 4 °C for at least 1 week before further analysis. SPB was prepared by mixing sodium phosphate dibasic anhydrous (NaH_2_PO_4_, cat. no. S375-500, Fisher Scientific) with sodium phosphate monobasic monohydrate (NaH_2_PO_4_•H_2_O, cat. no. S468-500, Fisher Scientific).

### Optical characterization

Three-dimensional excitation-emission matrices (EEM) were collected on a fluorometer (FluoroMax-4, Horiba) using a quartz cuvette (cat. no. 16.100F-Q-10/Z15, Sterna Cells). The scan ranges for both excitation and emission were set to be 400-800 nm in increments of 5 nm. The slit size and integration time were 5 nm and 0.1 s, respectively. Unless otherwise stated, 200 µl of 1 µM sample was used for one measurement. The acquired data were post-processed and visualized using a Python script.

All other fluorescence data were acquired using a plate reader (SpectraMax i3, Molecular Devices) and half-area UV transparent 96-well microplates (cat. no. 675801, Greiner Bio-One). For DNA/AgNCs, the excitation wavelength was set to 280 nm, while 490 nm was used for RNaseAlert. Each measurement was averaged over 15 lamp flashes, with the read height optimized prior to measurement. Each well contained 50 µl of sample. Emission spectra were scanned with a 5 nm increment, and data were collected every 2 minutes at 37 °C.

Absorbance data were also collected using the same plate reader and microplate. The absorption was scanned from 300 to 700 nm in increments of 5 nm. The concentration of all DNA-related samples was quantified and validated using a spectrophotometer (NanoDrop One Microvolume UV-Visible Spectrophotometer, ThermoFisher Scientific).

### Gel electrophoresis

Handcast polyacrylamide gels (20%) were prepared. For example, 16 ml of 20% gel was prepared as follows: 6.224 ml of nuclease-free water was added to 8 ml of 40% acrylamide solution (cat. no. HC2040), and the solution was mixed with 1.6 ml of 10× Tris-borate-ethylenediaminetetraacetic acid (TBE) buffer (cat. no. BP1333-1, Fisher Scientific). Then, 16 µl of N,N,N′,N′-tetramethylethylenediamine (TEMED) (cat. no. HC2006) was added. After mixing 160 µl of 10% ammonium persulfate (cat. no. HC2005), the gel solution was quickly poured into an empty gel cassette (cat. no. NC2010) installed in a customized handcast station. A gel comb (cat. nos. NC3002, NC3010 and NC3012) was then slowly inserted, and the gel was polymerized for >30 min at room temperature.

Gel electrophoresis was conducted using XCell SureLock Mini-Cell (cat. no. El0001) connected to a power supply (cat. no. PS0120). Sample (20 μl for 10 wells and 15 µl for 15 wells) was loaded into each well of a precast gel. The sample was loaded with a 10× TBE loading buffer, which was prepared by mixing 2 ml of 10× TBE buffer with 3 g Ficoll type 400 (cat. no. F10400-10.0, Research Products International) and nuclease-free water to 10 ml. The sample was concentrated up to 250 μM using a 3 kDa molecular weight cut-off (MWCO) centrifugal filter (cat. no. UFC5003, Sigma-Aldrich). Every gel contained at least one ladder lane using a DNA oligo standard (cat. no. 51-05-15-01, IDT) labelled with SYBR gold (cat. no. S11494, Thermo Fisher Scientific). Then, the gel was run at 140 V for 180 min in 1× TBE buffer unless stated otherwise and visualized using a UV transilluminator for true color images (model no. GV4L20X20, Syngene). The UV transilluminator is equipped with a 365 nm UV lamp, and the true color UV photos were taken by an iPhone 15 Pro (Apple) using Adobe Lightroom (ISO 500, 1/40 s for shutter speed and f/1.6 aperture) in a custom-built imaging system^1^. Materials for gel electrophoresis were purchased from Thermo Fisher Scientific unless stated otherwise.

### Gel purification

Subak substrates were purified by a modified crush-and-soak method using high-percentage PAGE. Then, 350 µl of 150 µM Subak substrates were loaded into a 20% handcast PAGE gels with a two-dimensional well comb. The loading sample included 35 µl of the aforementioned 10× loading buffer. After running at 140 V for 180 min, the gel was visualized under a UV transilluminator, and then a fluorescent gel slab was cut and eluted using a gel cutter (cat. no. 10048-876, VWR). This eluted gel was crushed as small as possible and then soaked in 20 mM SPB (pH 7.4) overnight. This solution was transferred to a 0.45 µm polyvinylidene difluoride column filter (cat. no. F2517-6, Thermo Fisher Scientific), and then centrifuged at 7,000 g for 30 min to remove any remaining gel fragments. The concentrate was quantified by measuring A by a spectrophotometer and stored at 4 °C for future use otherwise used immediately.

### RNase A/T1 treatment

RNase A/T1 Mix (cat. no. EN0551, Thermo Fisher Scientific) was used to evaluate the cleavage performance of rSubak under increasing nuclease concentrations. This commercial enzyme mixture contains 2 mg/ml of RNase A and 5000 U/ml of RNase T1, combining cytidine/uridine-and guanosine-specific ribonuclease activity in a single formulation. To determine the analytical performance of the substrate, we prepared a series of reactions with final RNase concentrations ranging from 0 to 10,000 pg/µl (0-0.5 µl enzyme per 50 µl reaction volume). All reactions were conducted in a 20 mM sodium phosphate buffer (SPB, pH 7.4), varying volumes of enzyme mix, and rSubak substrates at 5 μM final concentration. Nuclease-free water was used to adjust the final volume to 50 µl.

Reaction mixtures were assembled in UV-transparent 96-well plates and briefly centrifuged. The plates were incubated at 37 °C and monitored for green-to-red fluorescence conversion using kinetic fluorescence measurements, with data collected every 20 seconds over 60 minutes. Emission intensities at 530 nm and 625 nm were recorded using a plate reader, and ratiometric signal (I_625_/I_530_) was used to assess the extent of enzymatic cleavage.

### LoD calculation

Data were fitted using a four-parameter logistic (4-PL) sigmoidal model. The limit of detection (LoD) was calculated as the concentration corresponding to the mean signal of the blank control plus two standard deviations (2 S.D.). All data points are presented as the mean of duplicate measurements (n=2), with error bars representing the standard error of the mean (s.e.m.).

### MS

Prior to electrospray ionization (ESI)-MS, solutions containing 10 μM Subak substrates in 10 mM ammonium acetate were purified using Micro Bio-Spin P-6 Gel Columns (Bio-Rad). Octylamine was added to the solution at a concentration of approximately 0.1% (vol/vol) to reduce the adduction of alkali metals during the ESI process. The samples (diluted to 5 μM) were directly infused into an Orbitrap Fusion Lumos mass spectrometer (Thermo Fisher Scientific) using Au/Pd-coated borosilicate emitters fabricated in-house for nESI. Mass spectra were collected in negative-ionization mode over the range spanning *m/z* 400 and 3,000 using a resolving power of 120,000 at *m/z* 200, an automatic gain control target of 1E6 charges, and an ionization voltage of 550-600 V. All mass spectra were deconvoluted using Xtract in FreeStyle version 1.8 (Thermo Fisher Scientific). The following deconvolution parameters were applied: signal-to-noise threshold of 3, fit factor of 80%, and remainder threshold of 25%. Theoretical isotopic distributions were calculated using ChemDraw (Revvity Signals) and Xcalibur Qual Browser (Thermo Fisher Scientific). All mass spectra were interpreted and annotated manually with the aid of Mongo Oligo Mass Calculator v2.07. The mass spectrometer was calibrated using a mixture of components designed for the negative ionization mode to ensure high mass accuracy (typically ≤10 ppm error).

### In vitro transcription (IVT) for RNA virus synthesis

A total reaction volume of 100 µl was prepared by combining 75 µl deionized water (DIW), 10 µl DNase I reaction buffer, 10 µl of 100 µM template oligonucleotide and 15 µl of 100 µM primer. The solution was heated to 95 °C for 3 minutes to allow annealing and then gradually cooled to room temperature to enable dsDNA formation.

For IVT, a 30 µl reaction mixture was prepared in DNA LoBind PCR tubes. The mixture contained 6.25 µl DIW, 3 µl of 10× T7 RNA polymerase reaction buffer, 12 µl of 100 mM NTP mix (3 µl each of ATP, CTP, GTP, and UTP), 5 µl of 10 µM dsDNA template (>2 µg), 1.5 µl of 0.1 M dithiothreitol (DTT), and 2.25 µl of T7 RNA polymerase mix. The reaction was incubated at 37 °C for 16 hours. Following incubation, 3 µl of DNase I was added to the reaction to degrade the DNA template. The mixture was gently flicked, briefly centrifuged, and incubated at 37 °C for an additional 10-15 minutes. Lastly, the RNA product was purified using the Monarch RNA Cleanup Kit (NEB, Cat. #T2040), following the manufacturer’s protocol for <50 µg input.

### In vitro CRISPR/Cas13a reaction

Recombinant *Leptotrichia wadei* Cas13a (LwaCas13a) protein was purchased from GenScript, and rCutSmart buffer was obtained from New England Biolabs (cat. no. B6004S). Ribonucleoprotein (RNP) complexes were formed by combining 2.2 µl of Cas13a stock protein, 8.85 µl of crRNA (300 ng/µL), 10 µl of 10× rCutSmart buffer, and 78.95 µl of nuclease-free water to a final volume of 100 µL. This mixture yielded final concentrations of 125 nM Cas13a and 250 nM crRNA, based on an optimized 1:2 ratio. The RNP complex was incubated at 37 °C for 10 minutes to ensure full complexation prior to use.

For the CRISPR/Cas13a assay, 50 µl reactions were assembled containing 10 µl of RNP, 2.5 µl of 10 µM rSubak substrate (final 500 nM), 5 µl of 10× rCutSmart buffer, 5 µl of 0.7% Ficoll 400, variable volumes of synthetic RNA targets corresponding to final concentrations of 0-100 nM (0-100,000 pM), and nuclease-free water to complete the volume. Target concentrations tested included: 0, 0.5, 2, 10, 50, 200, 1000, 10,000, 50,000, and 100,000 pM. A no-target control (NTC) was included in each experimental run.

In parallel, a comparative assay was conducted using RNaseAlert (IDT, cat. no. 11-04-02-04), in which 12.5 µl of 2 µM RNaseAlert stock was added to each 50 µl reaction to achieve a final concentration of 500 nM. Reactions were incubated at 37 °C and monitored in real time using a plate reader (SpectraMax i3, Molecular Devices), with fluorescence recorded every 2 minutes over 30 minutes. Emission intensities at 530 nm (green) and 625 nm (red) were recorded for rSubak, while RNaseAlert was monitored using its standard green channel (520 nm). Each virus assay (SARS-CoV-2, influenza A (A/H5N1), and measles viral strands) was performed in duplicate per condition using rSubak-2 and RNaseAlert. To quantify the color-switching response, fluorescence intensity ratios (I_625_/I_530_) were calculated. These ratios represent the quotient of the red emission (I_625_)_and green emission (I_530_) signals measured after a 30-minute incubation in the presence of the target.

### Human serum sample preparation

Normal human serum (Invitrogen, cat. no. 31876) was used to evaluate matrix effects in CRISPR/Cas13a assays. The lyophilized serum was reconstituted according to the manufacturer’s instructions by adding 2.0 mL of nuclease-free distilled water, yielding a final concentration of 60 mg/ml total protein. The reconstituted serum was gently mixed and left at room temperature for 1-2 hours to fully dissolve. Prior to use in assays, the serum was heat-inactivated at 95 °C for 10 minutes to denature endogenous enzymes and was then centrifuged at 16,500 × g for 20 minutes^34,36^. The resulting supernatant was collected and either used immediately or stored in aliquots at −20 °C for future experiments.

For CRISPR/Cas13a reactions, the inactivated serum of a desired concentration (0.5-10%) was added to the reaction mixture, replacing an equivalent volume of nuclease-free water. All other reagent concentrations, including rCutSmart buffer, crowding agents, RNP complexes, and fluorescent substrates, were held constant. This setup was used to assess rSubak-2 performance under biologically relevant conditions and to evaluate its robustness in the presence of complex sample matrices.

### Secondary structure prediction and molecular visualization

Secondary structure folding and thermodynamic parameters (ΔG_folding_ and T_m_) for the DNA oligonucleotides were predicted using the UNAFold algorithm via the IDT SciTools (Integrated DNA Technologies). The folding simulations were performed under the following environmental assumptions to mimic experimental conditions: 25 °C, 50 mM Na^+^, 3 mM Mg^2+^, and a strand concentration of 0.2 µM.

To visually illustrate the proposed dual-path mechanism and the conformational states of the rSubak scaffolds, secondary structures were predicted using RNAfold via the ViennaRNA Web Services. These 2D predictions provided the basis for generating representative 3D architectures through RNAComposer. All final molecular renderings and high-resolution visualizations of the DNA/RNA-AgNC complexes were performed using PyMOL to represent the structural transitions and cluster-host interactions described in this study.

## Data availability

## Code availability

## Acknowledgements

This work was supported by the National Science Foundation grants (CBET2029266 to J.S.B. and 2404334 to H.-C.Y.) and the National Institutes of Health grant (EY033106 to H.-C.Y.). J.S.B is also gracious to the support of Welch Foundation (F1155).

## Author contributions

S.K., S.H. and H.-C.Y. conceived the project and designed the experiments. S.H. originally designed the template sequences for Subak substrates and mutation tests to optimize the substrates. S.K. performed the majority of experiments and proposed the mechanism of digestion-induced AgNC transformation. S.K., S.H. and J.N.W. prepared the samples for MS measurements. J.N.W. performed MS measurements and data analysis with assistance from J.S.B. J.S.B. supervised all MS experiments and data analysis. Y.H. assisted the curve fitting. A.-T.N. checked the buffer compatibility for RNase A/T1 digestion and in vitro CRISPR/Cas13a reaction. W.-R.C. and S.S. assisted the gel purification. Y.-A.K. and Y.-I.C. reviewed and edited the manuscript. S.K., S.H. and H.-C.Y. wrote the article with input from all authors. S.H. and H.-C.Y. supervised the project.

## Additional information

