## Supplementary information for "Mechanistic insights into the color transformation of a non-FRET substrate for RNase activity detection"

**Supplementary Table 1. List of sequences for the original Subak and rSubak substrates, initially tested based on the original motifs.**

(When a sequence is a hairpin DNA, the overhang arms, stem, and loop are colored in orange, blue, and black, respectively.)

| **Sequence name** | **Sequence (5'->3')** |
|---|---|
| Subak-1 | AACCACCCCA TCGTCTT TTTT AAGACGA GTTCCCCGC |
| Subak-2 | AACCACCCCA TCGTCTC TTTT GAGACGA GTTCCCCCC |
| rSubak-1<br>(Subak-2_C17rC/C26rC) | AACCACCCCA TCGTCTrC TTTT GAGArCGA GTTCCCCCC |
| rSubak-2<br>(Subak-2_T16rU/C17rC<br>/T18-21rU) | AACCACCCCA TCGTCrUrC rUrUrUrU GAGACGA<br>GTTCCCCCC |
| pcSubak | AACCACCCCA TCGTCTC TTTT GAGACGA GTTCCCCCC |
| Subak-1_T16rU | AACCACCCCA TCGTCrUT TTTT AAGACGA GTTCCCCGC |
| Subak-1_T16rU/C26rC | AACCACCCCA TCGTCrUT TTTT AAGArCGA GTTCCCCGC |
| Subak-2_C17rC | AACCACCCCA TCGTCTrC TTTT GAGACGA GTTCCCCCC |

**Supplementary Table 2. Cost breakdown concerning rSubak substrate compared to FRET substrate.**

| | rSubak (ours) | Gel-purified rSubak (ours) | RNaseAlert (commercial) |
|---|---|---|---|
| DNA (from IDT) | $40.02 (250 nmol) | $40.02 (250 nmol) | |
| Silver nitrate ($485.00/50g from Sigma Aldrich) | | <$0.01 | |
| Sodium borohydride ($99.10/25g from Sigma Aldrich) | <$0.01 | <$0.01 | |
| Gel cassette ($230/25 units from Thermo Fisher) | n/a | $27.60 (3 gels for 210 nmol) | |
| Gel materials | n/a | <$0.01 | |
| 3K centrifugal filter ($518/96units from Sigma Aldrich) | n/a | $16.20 (3 filters) | |
| PVDF centrifugal filter ($234.53/100units from Thermo Fisher) | n/a | $7.05 (3 filters) | |
| Yield rate* | ~40% | ~40% | |
| Total cost (for 80 nmol) | <$40.04 | <$90.9 | |
| Total cost (per nmol) | <$0.50 | <$1.14 | $76 |

**Supplementary Table 3. List of sequences for modified rSubak substrates with varying number of consecutive ribonucleotides and local bulge motifs.**

(When a sequence is a hairpin DNA, the overhang arms, stem, loop, and bulge are colored in orange, blue, black, and red, respectively.)

| **No.** | **Sequence name** | **Sequence (5’->3’)** |
|---|---|---|
| R1 | Subak-2_bulge1 | AACCACCCCA TCGTCTrUrUC TTTT GAGACGA GTTCCCCCC |
| R2 | Subak-2_bulge2 | AACCACCCCA TCGTCTrUrUrUrUC TTTT GAGACGA GTTCCCCCC |
| R3 | Subak-2_bulge3 | AACCACCCCA TCGTCTrGrUC TTTT GAGACGA GTTCCCCCC |
| R4 | Subak-2_bulge4 | AACCACCCCA TCGTCTrGrUrGrUC TTTT GAGACGA GTTCCCCCC |
| R5 | Subak-2_T16rU/C17rC/T18-21rU (rSubak-2) | AACCACCCCA TCGTCrUrC rUrUrUrU GAGACGA GTTCCCCCC |
| R6 | Subak-2_C17rC/T18-21rU | AACCACCCCA TCGTCTrC rUrUrUrU GAGACGA GTTCCCCCC |
| R7 | Subak-2_T16rU/C17rC/T18-21rU/C26rC/G27rG/A28rA | AACCACCCCA TCGTCrUrC rUrUrUrU GAGArCrGrA GTTCCCCCC |
| R8 | Subak-2_ T16rU/C17rC/T18-21rU/G27rG/A28rA | AACCACCCCA TCGTCrUrC rUrUrUrU GAGACrGrA GTTCCCCCC |

**Supplementary Table 4. List of sequences for CRISPR/Cas13a reaction.**

| | Sequence name | Sequence (5'->3') | Size (nt) | Ref. |
|---|---|---|---|---|
| crRNA (Lwa Cas13a) | SARS-CoV-2 | /AltR1/rGrGrGrArUrUrUrArGrArCrUrArCrCrCrCrArArArArArCrGrArArGrGrGrGrArCrUrArArArArCrGrCrArGrCrArCrCrArGrCrUrGrUrCrCrArArCrCrUrGrArArGrArArG/AltR2/ | 61 | 1 |
| | Measles virus (MV) | /AltR1/rGrGrGrArUrUrUrArGrArCrUrArCrCrCrCrArArArArArCrGrArArGrGrGrGrArCrUrArArArArCrArGrCrUrCrArUrUrCrUrCrArCrUrUrGrArUrCrArCrCrGrUrGrUrA/AltR2/ | 61 | 2 |
| | Influenza A (A/H5N1) | /AltR1/rGrGrGrArUrUrUrArGrArCrUrArCrCrCrCrArArArArArCrGrArArGrGrGrGrArCrUrArArArArCrCrUrGrArArCrArGrUrGrUrCrArArArUrGrUrUrCrCrArArGrCrArCrA/AltR2/ | 61 | 3 |
| RNA viral fragment | T7_SARS-CoV-2 | GAATTAATACGACTCACTATAGGACTCCTGGTGATTCTTCTTCAGGTTGGACAGCTGGTGCTGCAGCTTATTATGTGGGTTATCTTCAACCTAGGACTTT | 100 | 1 |
| | 0T7_SARS-CoV-2 | AAAGTCCTAGGTTGAAGATAACCCACATAATAAGCTGCAGCACCAGCTGTCCAACCTGAAGAAGAATCACCAGGAGTCCTATAGTGAGTCGTATTAATTC | | |
| | T7_MV | GAATTAATACGACTCACTATAGGTCCTGGAAGAACAAAGTTCAGACACGGACACCCCCAGGGTGTACAATGACAGAGATCTTCTAGAC | 88 | 2 |
| | 0T7_MV | GTCTAGAAGATCTCTGTCATTGTACACCCTGGGGGTGTCCGTGTCTGAACTTTGTTCTTCCAGGACCTATAGTGAGTCGTATTAATTC | | |
| | T7_A/H5N1 | GAATTAATACGACTCACTATAGGAATGCGAGATGTGCTTGGAACATTTGACACTGTTCAGATAATAAAACTTCTCCCCTTTGCTGCTGCCCCA | 93 | 3 |
| | 0T7_A/H5N1 | TGGGGCAGCAGCAAAGGGGAGAAGTTTTATTATCTGAACAGTGTCAAATGTTCCAAGCACATCTCGCATTCCTATAGTGAGTCGTATTAATTC | | |

[*]Alt-R1/Alt-R2 denote proprietary chemical modifications (2′-O-methyl and phosphorothioate linkages) provided by IDT to increase gRNA stability and potency.

[1]https://go.idtdna.com/rs/400-UEU-432/images/Zhang%20et%20al.%2C%202020%20COVID-19%20detection%20(updated).pdf

[2]https://journals.asm.org/doi/10.1128/jcm.01456-24 (Zubach et al., J. Clin. Microbiol., 2025)

[3]https://pubmed.ncbi.nlm.nih.gov/39387583/ (Timarco et al., J. Virol., 2024)

**Supplementary Table 5. List of sequences for modified rSubak substrates based on Subak-1 motif.** First 3 rows have 1 cleavage site (cleaving only 5'-side) and other 7 rows have 2 cleavage sites (cleaving both 5'-side and 3'-side).
(When a sequence is a hairpin DNA, the overhang arms, stem, loop, and bulge are colored in orange, blue, black, and red, respectively.)

| No. | Sequence name | Sequence (5'->3') |
|---|---|---|
| V1 | Subak-1_T16rU/T17rU | AACCACCCCA TCGTCrUrU TTTT AAGACGA GTTCCCCGC |
| V2 | Subak-1_T16-21rU | AACCACCCCA TCGTCrUrU rUrUrUrU AAGACGA GTTCCCCGC |
| V3 | Subak-1_T25rU/T26rU | AACCACCCCA TCGTCTT TTTT AAGrUrUACGA GTTCCCCGC |
| V4 | Subak-1_T17rU/T18rU/T27rU/T28rU | AACCACCCCA TCGTCTrUrUT TTTT AAGrUrUACGA GTTCCCCGC |
| V5 | Subak-1_T17rU/T18rU/C28rC | AACCACCCCA TCGTCTrUrUT TTTT AAGArCGA GTTCCCCGC |
| V6 | Subak-1_T16rU/C26rC | AACCACCCCA TCGTCrUT TTTT AAGArCGA GTTCCCCGC |
| V7 | Subak-1_T17rU/C26rC | AACCACCCCA TCGTCTrU TTTT AAGArCGA GTTCCCCGC |
| V8 | Subak-1_T16rU/T25rU/T26rU | AACCACCCCA TCGTCrUT TTTT AAGrUrUACGA GTTCCCCGC |
| V9 | Subak-1_T16rU/T17rU/T25rU/T26rU | AACCACCCCA TCGTCrUrU TTTT AAGrUrUACGA GTTCCCCGC |
| V10 | Subak-1_T16rU/T17rU/T25-28rU | AACCACCCCA TCGTCrUrU TTTT AAGrUrUrUrUACGA GTTCCCCGC |

**Supplementary Table 6. Limit of detection (LoD) for three RNA viruses using CRISPR/Cas13a in sodium phosphate buffer (SPB, pH 7.4) and 2% human serum.**

| | | Buffer (SPB, pH 7.4) | | | 2% human serum | | |
|---|---|---|---|---|---|---|---|
| Parameter | | Subak ($I_{625}$) | Subak ($I_{625}/I_{530}$) | RNaseAlert ($I_{520}$) | Subak ($I_{625}$) | Subak ($I_{625}/I_{530}$) | RNaseAlert ($I_{520}$) |
| LoD ($\mu + 3\sigma$, pM) | SARS-CoV-2 | 2.6 | 0.7 | 130 | 21 | 13 | 31 |
| | A/H5N1 | 0.6 | 0.3 | 248 | 0.8 | 0.4 | 253 |
| | MV | 54 | 57 | 152 | 1.6 | 16 | 1,047 |

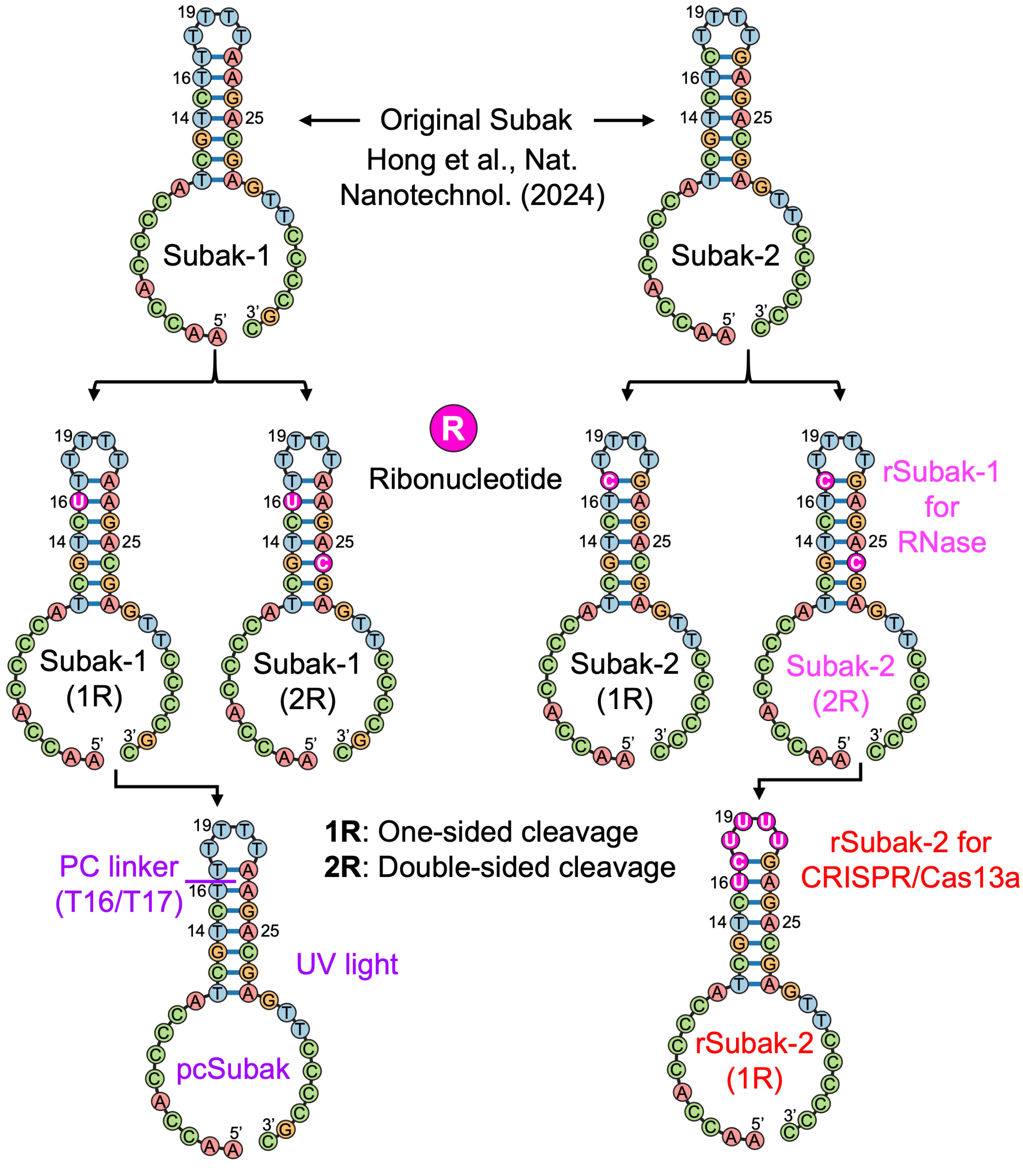

**Supplementary Fig. 1. Design and structural evolution of the Subak substrate family.** Schematic diagrams illustrating the systematic transition from the two original DNA-based Subak substrates (Subak-1 and -2) to various variants. This evolution includes the rSubak variants with one- (1R) or double (2R) ribonucleotide cleavage sites, specifically rSubak-1 for RNase detection and rSubak-2 for CRISPR/Cas13a, as well as the pcSubak (photocleavable) variant. R denotes the distribution of ribonucleotide cleavage sites: 1R indicates substitutions on a single-sided stem (left or right), while 2R signifies substitutions on both stems, regardless of the total number of nucleotides.

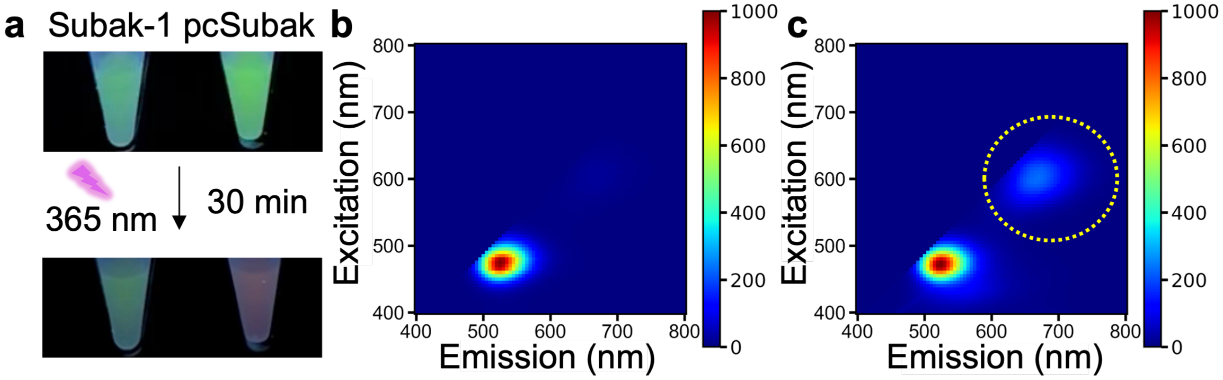

**Supplementary Fig. 2. Characterization of UV-activated color transformation in pcSubak.** Experimental validation of the photocleavable Subak (pcSubak) system for light-induced color-change. (**a**) Fluorescence color of Subak-1 and pcSubak before and after 30 min of UV irradiation (365 nm) using a UV transilluminator. Only pcSubak exhibited a green-to-red color shift. (**b, c**) Excitation-emission matrix (EEM) spectra of pcSubak before (**b**) and after (**c**) 30 min of UV irradiation, demonstrating a transition to red fluorescence. Further optimization is required to enhance the color shift efficiency.

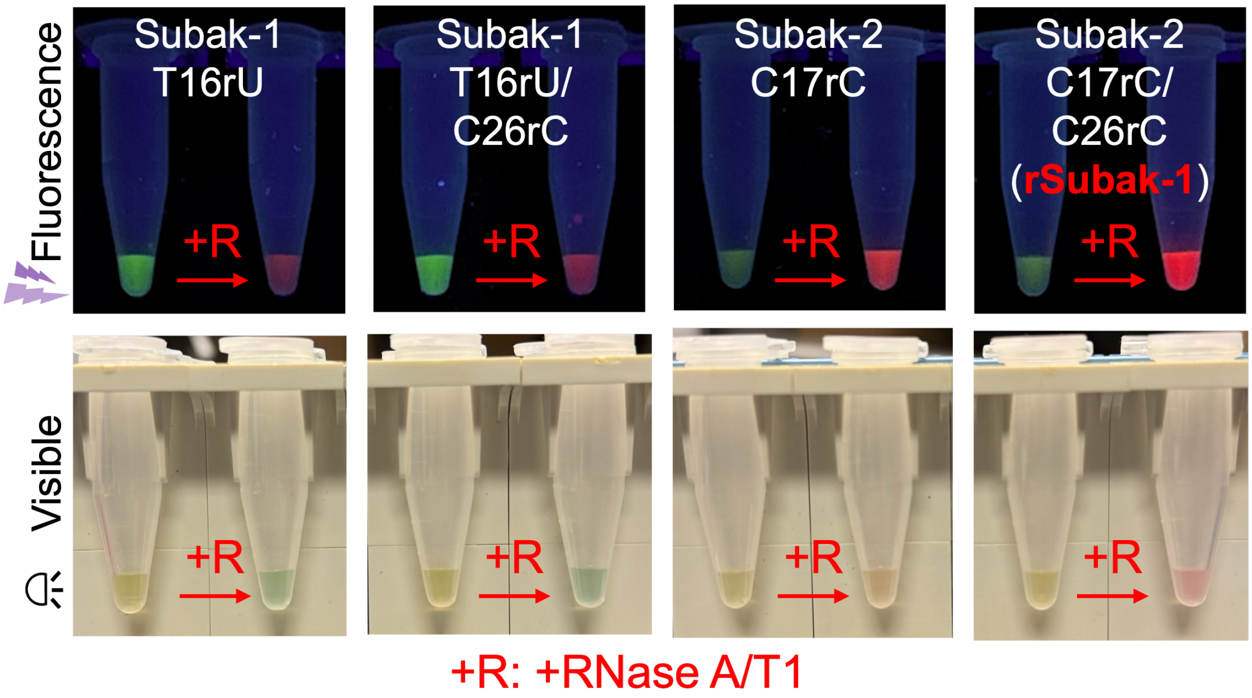

**Supplementary Fig. 3. Fluorescence and colorimetric images of rSubak substrates before and after RNase A/T1 digestion.** Photographic characterization of Subak-1 and Subak-2 variants demonstrating distinct color changes upon RNase A/T1 cleavage. Under UV illumination (top row), all variants exhibit a clear transition from initial green fluorescence (left tube) to red emission (right tube) upon cleavage, while rSubak-1 showing the most significant change. Under visible light (bottom row), the substrates undergo a subtle yet distinct color change from pale yellow to green, orange, or pinkish-red. These results demonstrate that rSubak substrates can also serve as colorimetric sensors for point-of-care (POC) diagnostics.

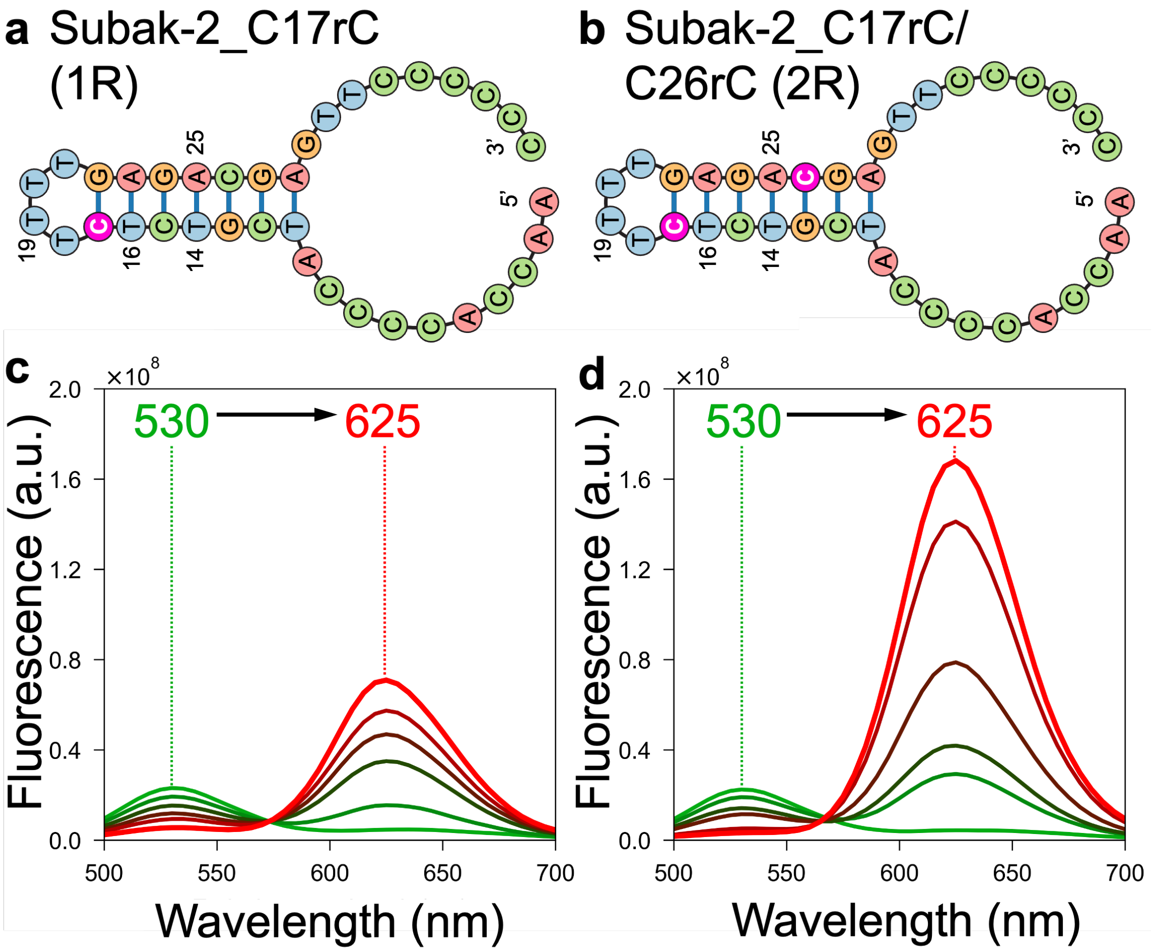

**Supplementary Fig. 4. Concentration-dependent fluorescence spectra (0-4000 pg/µl) of Subak-2 variants.** (**a,b**) Schematic structures of the Subak-2 1R (**a**) and 2R (**b**, rSubak-1) variants. (**c,d**) Fluorescence emission spectra of the 1R (**c**) and 2R variant (**d**, rSubak-1) in response to increasing concentrations of RNase A/T1 (0-4000 pg/µl). Both variants exhibit a distinct green-to-red conversion (95 nm shift). While the 1R variant shows no significant residual green emission here, the 2R variant (**d**, rSubak-1) achieves substantially higher red emission, demonstrating that double site cleavage enhances the overall signal-on response. R denotes the distribution of ribonucleotide cleavage sites: 1R indicates substitutions on a single stem (left or right), while 2R signifies substitutions on both stems, regardless of the total number of nucleotides.

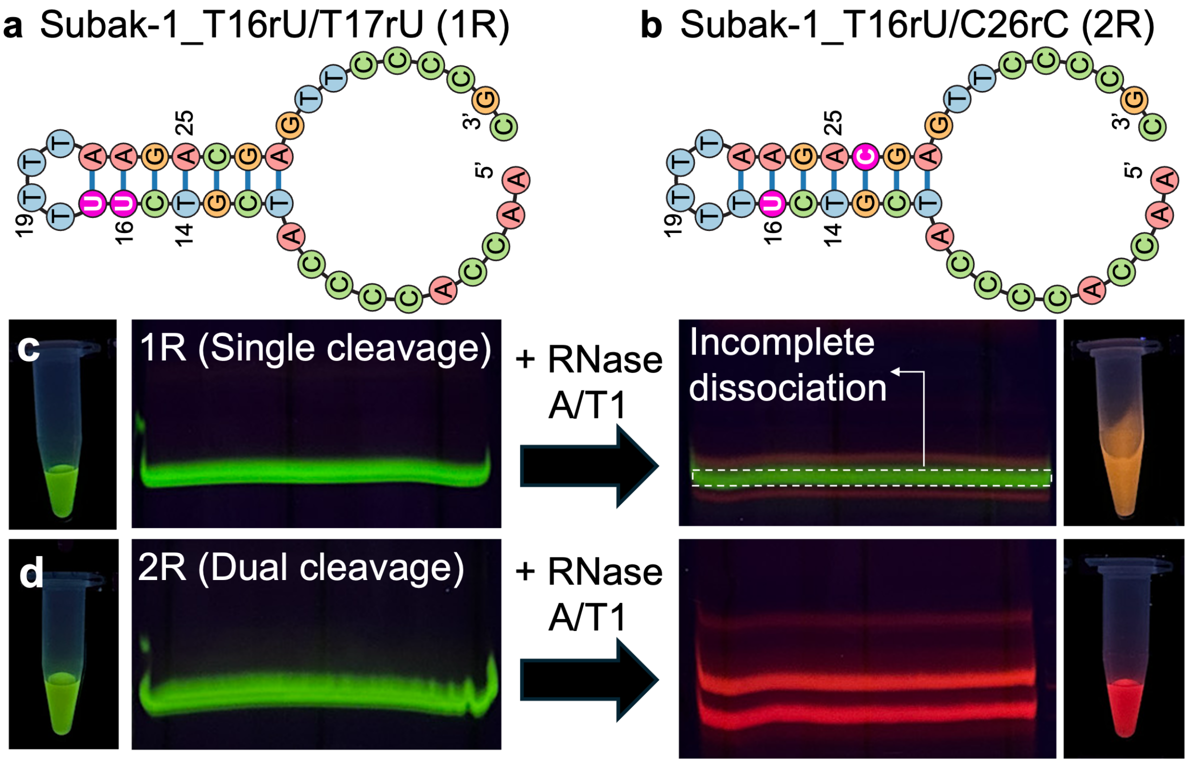

**Supplementary Fig. 5. PAGE analysis of green-to-red transformation in Subak-1 variants.** (**a,b**) Schematic structures of 1R (one-sided cleavage) and 2R (double-sided cleavage) variants. Ribonucleotides are colored in magenta. (**c,d**) Fluorescence images of tubes and gel bands for 1R (**c**) and 2R (**d**) substrates before and after RNase A/T1 addition. While both substrates are initially green, only the 2R variant (Subak-1_T16rU/C26rC) shows a complete shift to red fluorescence and distinct red gel bands after cleavage. The 1R variant retains residual green-emitting species. Tube photographs were captured under UV illumination (365 nm). PAGE was performed using a 20% gel at 140 V for 3 hours and imaged under the same excitation source as the tube imaging.

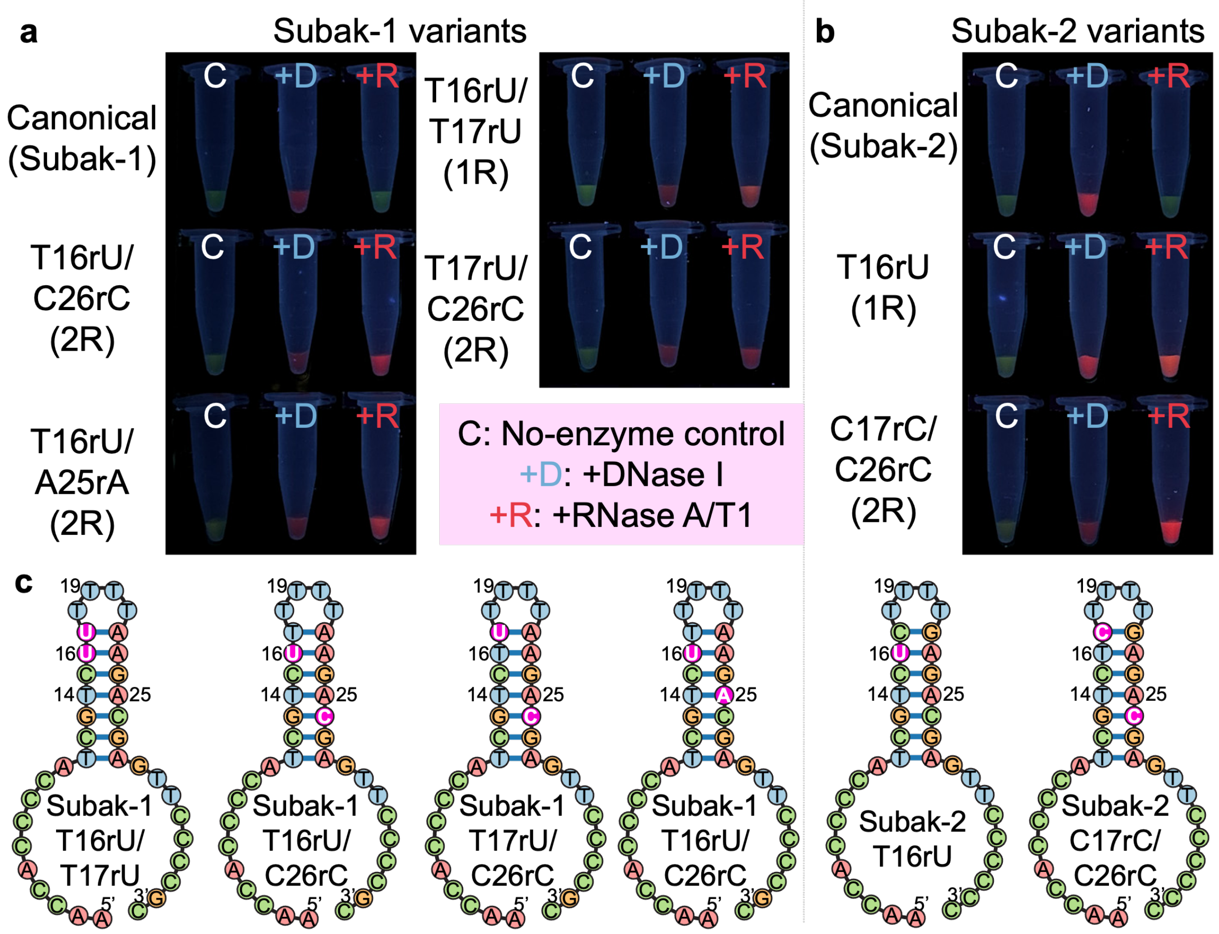

**Supplementary Fig. 6. Characterization of color change in one- and double-sided cleavage rSubak variants.** Fluorescence images (under 365 nm excitation) of Subak-1 (**a**) and Subak-2 (**b**) variants with either one (1R) or two (2R) ribonucleotide cleavage sites. The 1R design denotes a single ribonucleotide substitution in the left stem (e.g., T16rU or T17rU), while 2R denotes substitutions at two positions in both stem regions (e.g., T16rU/C26rC). For all variants, no-enzyme controls (C) retain their initial green fluorescence. Non-specific DNase I treatments (+D) show minimal red emission, whereas digestion with RNase A/T1 (+R) triggers a robust color change. For both variants, the 2R constructs achieve a more complete green-to-red spectral conversion and higher red fluorescence intensity compared to 1R constructs. Reactions were performed with 10 µM substrate and 10 U enzyme at 37 °C for 30 min. (**c**) Schematic illustrations of the rSubak variant structures, where ribonucleotides are colored in magenta.

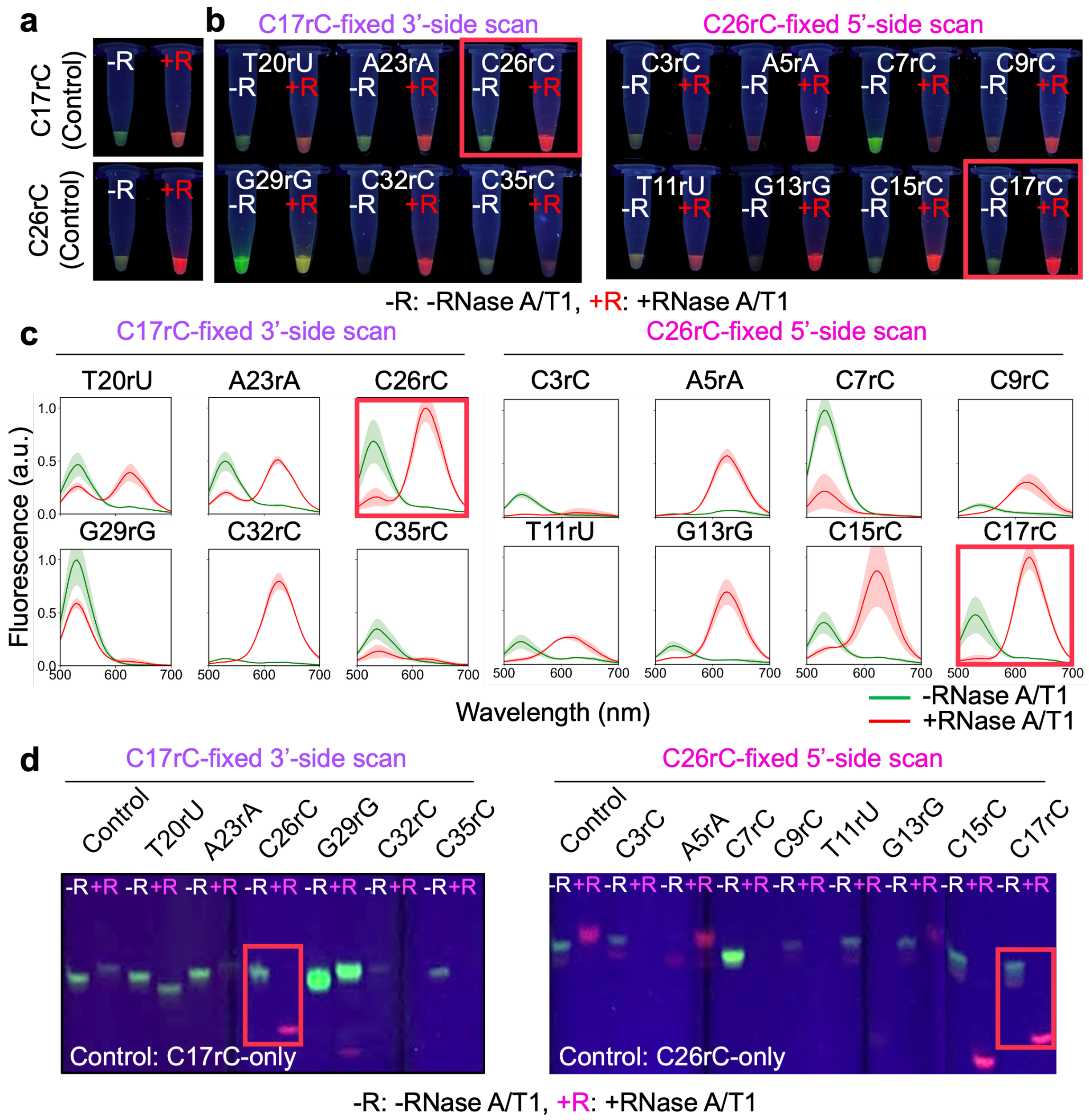

**Supplementary Fig. 7. Scanning and characterization of double-sided cleavage constructs.** Fluorescence images of (**a**) C17rC-only and C26rC-only controls were captured under 365 nm UV excitation before (-R) and after (+R) RNase A/T1 treatment. (**b**) Fluorescence images of the C17rC-fixed 3'-side substitution library and the C26rC-fixed 5'-side substitution library under identical UV illumination. (**c**) Emission spectra of the screened variants. All spectra were acquired under 280 nm excitation. For comparative visualization, intensities are normalized to the highest value within each group, except for the G29rG and C7rC variants; due to their exceptionally high emission, these two were normalized to their respective local maxima. (**d**) PAGE analysis confirms the enzymatic digestion across all variants. The C17rC/C26rC construct exhibits the most distinct and robust green-to-red color conversion, consistent with the tube images and spectra. Gel images were acquired using the same UV transillumination settings as the tube photographs in **a** and **b**.

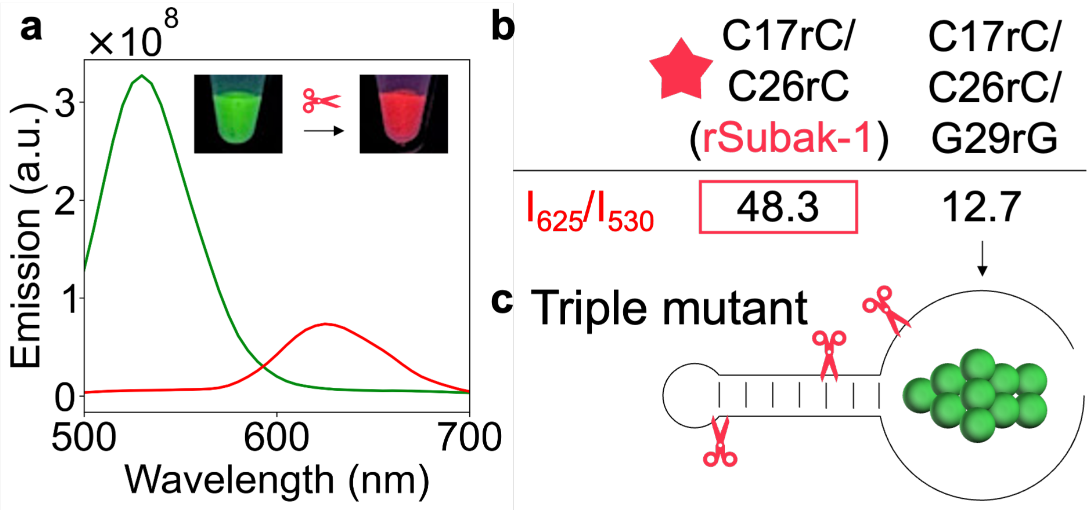

**Supplementary Fig. 8. Performance evaluation of triple-substitution rSubak substrate.** (**a**) Emission spectra of the C17rC/C26rC/G29rG triple mutant before (green) and after (red) RNase A/T1 cleavage. (**b**) Comparison of the ratiometric fluorescence intensity ($I_{625}/I_{530}$) between the optimal rSubak-1 (C17rC/C26rC) and the triple mutant construct. **(c**) Schematic representation of the triple mutant construct with three cleavage sites.

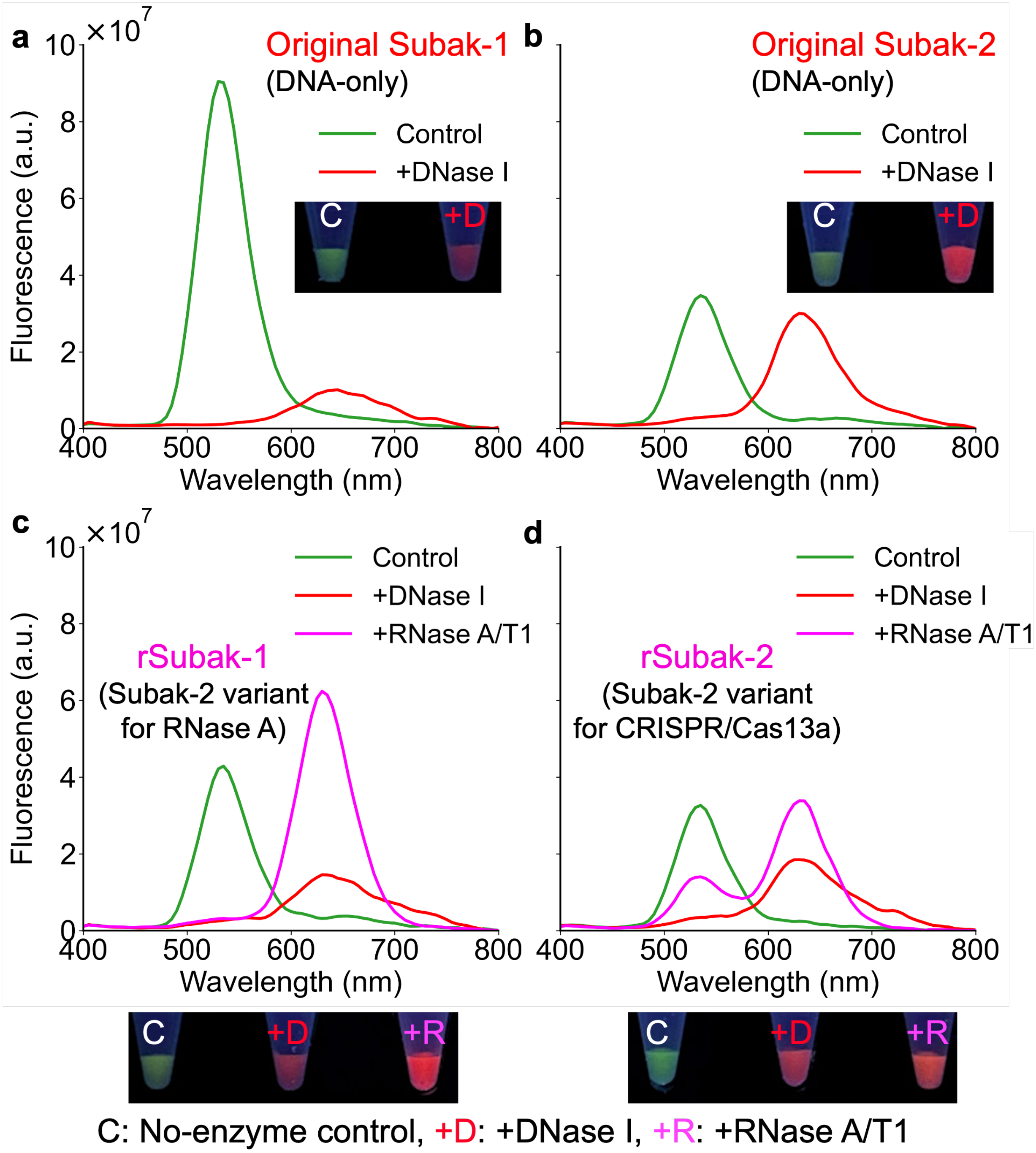

**Supplementary Fig. 9. Comparison of nuclease-induced cleavage between original Subak and optimized rSubak variants.** (**a-d**) Fluorescence emission spectra and corresponding photographs of DNA-only Subak-1 (**a**), Subak-2 (**b**), rSubak-1 (**c**), and rSubak-2 (**d**) before (C, green line) and after cleavage by DNase I (+D, red line) or RNase A/T1 (+R, magenta line). The rSubak variants, particularly rSubak-1, exhibit a robust color shift upon RNase A/T1 addition. Photographs were captured under UV illumination (365 nm), and fluorescence spectra were acquired using an excitation wavelength of 280 nm.

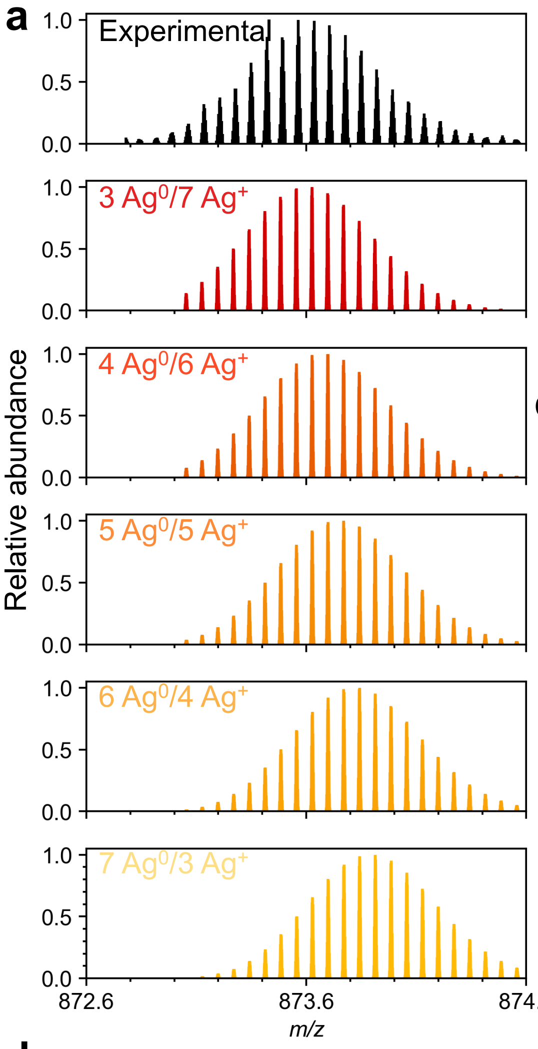

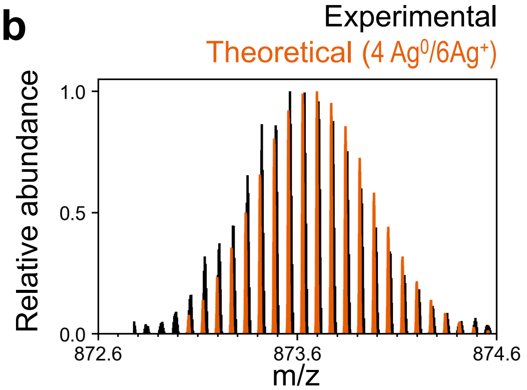

c

| Experimental *m/z* | Theoretical *m/z* | Mass Error *ppm* |
|---|---|---|
| 872.992512 | 872.983049 | 10.84 |
| 873.066247 | 873.054558 | 13.39 |
| 873.135266 | 873.126073 | 10.53 |
| 873.208089 | 873.197594 | 12.02 |
| 873.276746 | 873.269121 | 8.73 |
| 873.349196 | 873.340653 | 9.78 |
| 873.421993 | 873.411821 | 11.65 |
| 873.490841 | 873.483365 | 8.56 |
| 873.563151 | 873.554915 | 9.43 |
| 873.636118 | 873.626471 | 11.04 |
| 873.705409 | 873.698033 | 8.44 |
| 873.778157 | 873.76923 | 10.22 |
| 873.850818 | 873.840803 | 11.46 |
| 873.920063 | 873.912382 | 8.79 |
| 873.993148 | 873.983968 | 10.5 |
| 874.063382 | 874.055188 | 9.37 |
| 874.136574 | 874.126785 | 11.2 |

d

3 $Ag^0$/7 $Ag^+$
$C_{354}H_{\underline{433}}N_{126}O_{224}P_{36}Ag_{10}$

4 $Ag^0$/6 $Ag^+$
$C_{354}H_{\underline{434}}N_{126}O_{224}P_{36}Ag_{10}$

5 $Ag^0$/5 $Ag^+$
$C_{354}H_{\underline{435}}N_{126}O_{224}P_{36}Ag_{10}$

6 $Ag^0$/4 $Ag^+$
$C_{354}H_{\underline{436}}N_{126}O_{224}P_{36}Ag_{10}$

7 $Ag^0$/3 $Ag^+$
$C_{354}H_{\underline{437}}N_{126}O_{224}P_{36}Ag_{10}$

**Net charge of 14-**

e

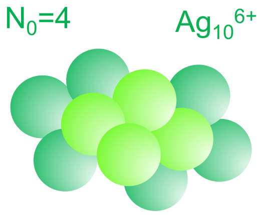

**Supplementary Fig. 10. Analysis of the isotopic distributions produced by ESI-MS of the intact rSubak-1 $Ag_{10}^{6+}$ green-emitting species.** (**a**) Comparison of the experimental and theoretical isotopic distributions for various $Ag^0$/$Ag^+$ ratios allows estimation of the stoichiometric composition of the intact rSubak-1 green cluster. (**b**) The experimental isotopic distribution aligns with the theoretical isotopic distribution for an $Ag_{10}$ cluster containing a ratio of 4 $Ag^0$/6 $Ag^+$. (**c**) The calculated mass error (ppm) values for the 4 $Ag^0$/6 $Ag^+$ cluster and the (**d**) chemical formulas of various $Ag^0$/$Ag^+$ ratios are presented. The net charge (14-) is balanced by the number of protons lost for each molecular composition. (**e**) The proposed structure of the 4 $Ag^0$/6 $Ag^+$ cluster is shown.

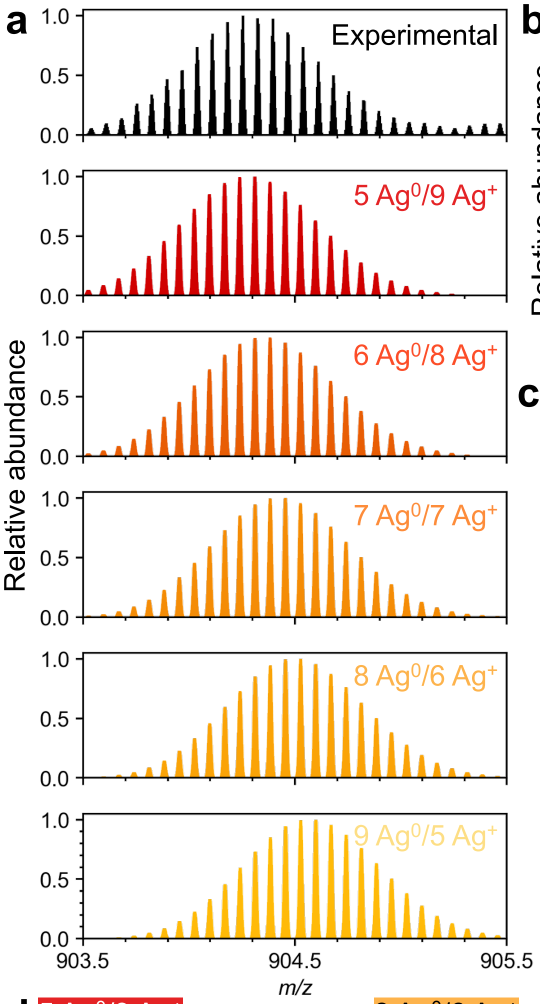

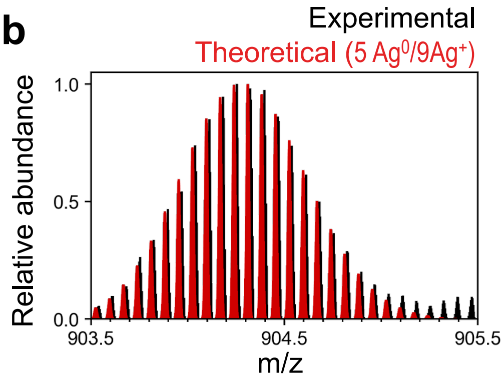

**c**

| Experimental *m/z* | Theoretical *m/z* | Mass Error *ppm* |
|---|---|---|
| 904.03864 | 904.024865 | 15.2 |
| 904.111675 | 904.096251 | 17.1 |
| 904.1812 | 904.168025 | 14.6 |
| 904.253911 | 904.239422 | 16.0 |
| 904.32331 | 904.310825 | 13.8 |
| 904.396338 | 904.382234 | 15.6 |
| 904.465761 | 904.453648 | 13.4 |
| 904.538513 | 904.525068 | 14.9 |
| 904.611862 | 904.596875 | 16.6 |
| 904.681043 | 904.668307 | 14.1 |
| 904.753948 | 904.739744 | 15.7 |
| 904.82387 | 904.811186 | 14.0 |
| 904.897036 | 904.882634 | 15.9 |
| 904.965824 | 904.954088 | 13.0 |
| 905.038957 | 905.025547 | 14.8 |
| 905.11223 | 905.097395 | 16.4 |
| 905.182757 | 905.168865 | 15.3 |

**d**

5 $Ag^0$/9 $Ag^+$: $C_{354}H_{\underline{431}}N_{126}O_{224}P_{36}Ag_{14}$

6 $Ag^0$/8 $Ag^+$: $C_{354}H_{\underline{432}}N_{126}O_{224}P_{36}Ag_{14}$

7 $Ag^0$/7 $Ag^+$: $C_{354}H_{\underline{433}}N_{126}O_{224}P_{36}Ag_{14}$

8 $Ag^0$/6 $Ag^+$: $C_{354}H_{\underline{434}}N_{126}O_{224}P_{36}Ag_{14}$

9 $Ag^0$/5 $Ag^+$: $C_{354}H_{\underline{435}}N_{126}O_{224}P_{36}Ag_{14}$

**Net charge of 14-**

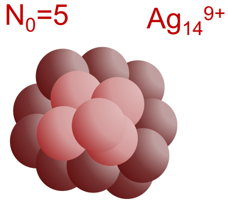

**Supplementary Fig. 11. Analysis of the isotopic distributions produced by ESI-MS of the intact rSubak-1 $Ag_{14}^{9+}$ dark-emissive species.** (**a**) Comparison of the experimental and theoretical isotopic distributions for various $Ag^0/Ag^+$ ratios allows estimation of the stoichiometric composition of the intact rSubak-1 dark cluster. (**b**) The experimental isotopic distribution aligns with the theoretical isotopic distribution for an $Ag_{14}$ cluster containing a ratio of 5 $Ag^0$/9 $Ag^+$ (**a, b**). (**c**) The calculated mass error (ppm) values for the 5 $Ag^0$/9 $Ag^+$ cluster and (**d**) the chemical formulas of various $Ag^0/Ag^+$ ratios are presented. The net charge (14-) is balanced by the number of protons lost for each molecular composition. (**e**) The proposed structure of the 5 $Ag^0$/9 $Ag^+$ cluster is presented.

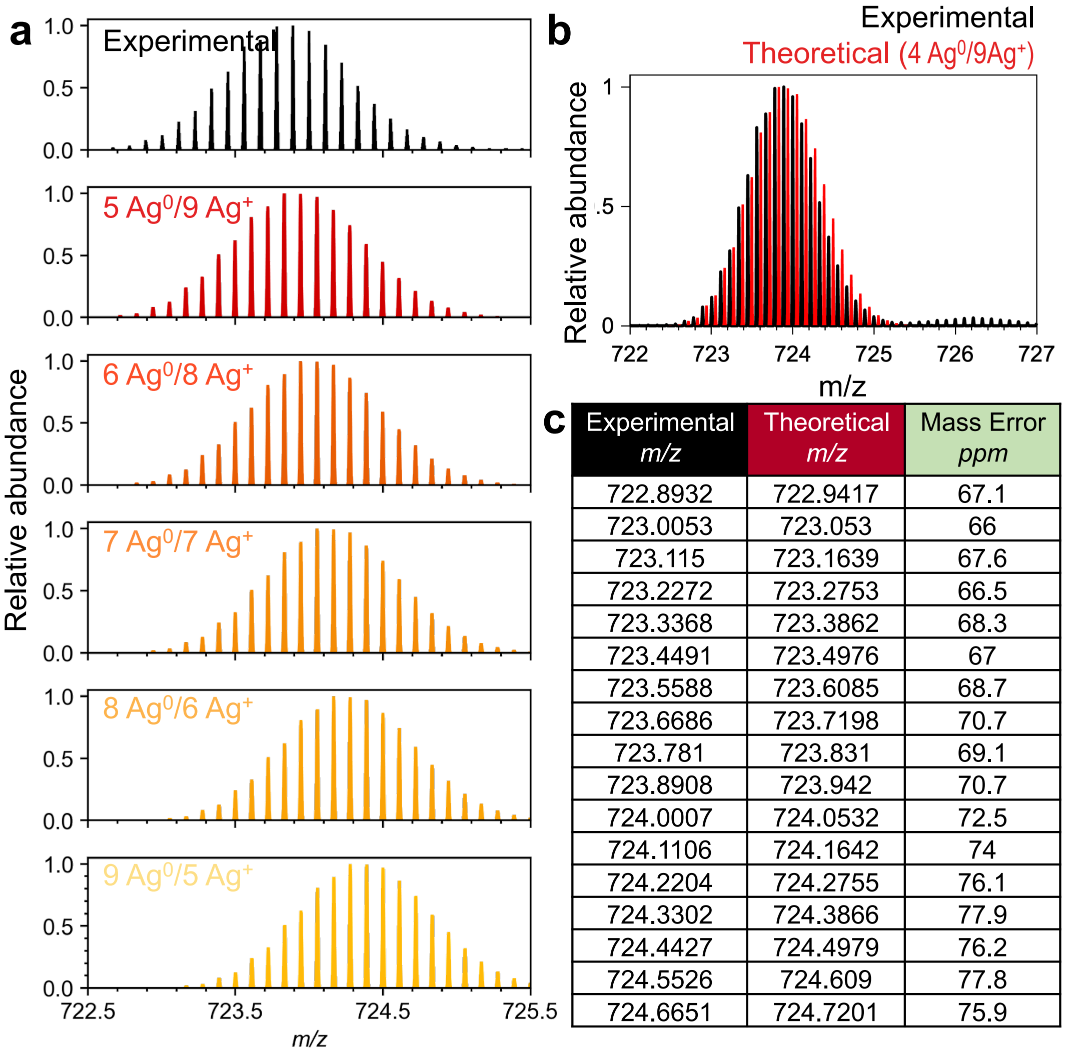

**c**

| Experimental *m/z* | Theoretical *m/z* | Mass Error *ppm* |
|---|---|---|
| 722.8932 | 722.9417 | 67.1 |
| 723.0053 | 723.053 | 66 |
| 723.115 | 723.1639 | 67.6 |
| 723.2272 | 723.2753 | 66.5 |
| 723.3368 | 723.3862 | 68.3 |
| 723.4491 | 723.4976 | 67 |
| 723.5588 | 723.6085 | 68.7 |
| 723.6686 | 723.7198 | 70.7 |
| 723.781 | 723.831 | 69.1 |
| 723.8908 | 723.942 | 70.7 |
| 724.0007 | 724.0532 | 72.5 |
| 724.1106 | 724.1642 | 74 |
| 724.2204 | 724.2755 | 76.1 |
| 724.3302 | 724.3866 | 77.9 |
| 724.4427 | 724.4979 | 76.2 |
| 724.5526 | 724.609 | 77.8 |
| 724.6651 | 724.7201 | 75.9 |

**d**

8 $Ag^0$/5 $Ag^+$
$C_{161}H_{191}N_{58}O_{103}P_{17}Ag_{13}$

7 $Ag^0$/6 $Ag^+$
$C_{161}H_{192}N_{58}O_{103}P_{17}Ag_{13}$

6 $Ag^0$/7 $Ag^+$
$C_{161}H_{193}N_{58}O_{103}P_{17}Ag_{13}$

5 $Ag^0$/8 $Ag^+$
$C_{161}H_{194}N_{58}O_{103}P_{17}Ag_{13}$

4 $Ag^0$/9 $Ag^+$
$C_{161}H_{195}N_{58}O_{103}P_{17}Ag_{13}$

**Net charge of 9-**

**e**

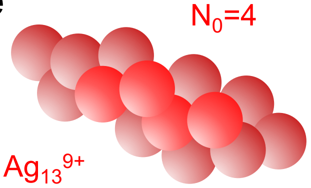

**Supplementary Fig. 12. Analysis of the isotopic distributions produced by ESI-MS of the digested rSubak-1 $Ag_{13}^{9+}$ red-emitting species.** (**a**) Comparison of the experimental and theoretical isotopic distributions for various $Ag^0/Ag^+$ ratios allows estimation of the specific cluster transformation associated with red fluorescence after enzymatic digestion. (**b**) The experimental isotopic distribution aligns with the theoretical isotopic distribution for an $Ag_{13}$ cluster containing a ratio of 4 $Ag^0$/9 $Ag^+$. (**c**) The calculated mass error (ppm) values for the 4 $Ag^0$/9 $Ag^+$ cluster and (**d**) the chemical formulas of various $Ag^0/Ag^+$ ratios are presented. The net charge (9-) is balanced by the number of protons lost for each molecular composition. (**e**) The proposed structure of the 4 $Ag^0$/9 $Ag^+$ cluster is presented, validating the signal-on state.

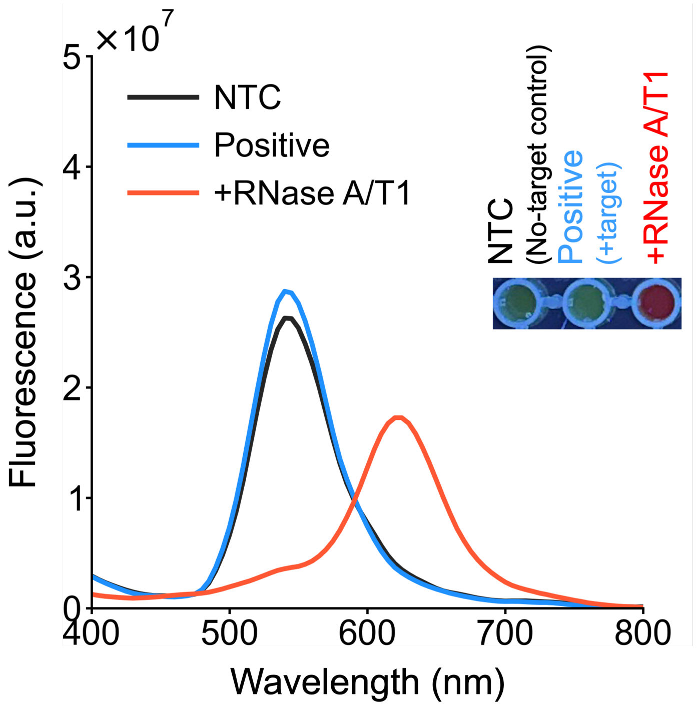

**Supplementary Fig. 13. Evaluation of rSubak-1 cleavage by CRISPR/Cas13a upon target recognition.** Fluorescence emission spectra of rSubak-1 (280 nm excitation) under different cleavage conditions. NTC (no-target control) represents rSubak-1 incubated with the LwaCas13a-crRNA complex (RNP) alone. Positive indicates the reaction containing both the RNP and its specific SARS-CoV-2 target. RNase A/T1 serves as a functional control for complete substrate digestion. While rSubak-1 exhibits a distinct green-to-red spectral shift upon RNase A/T1 cleavage, no discernible red emission is observed following Cas13a-mediated cleavage.

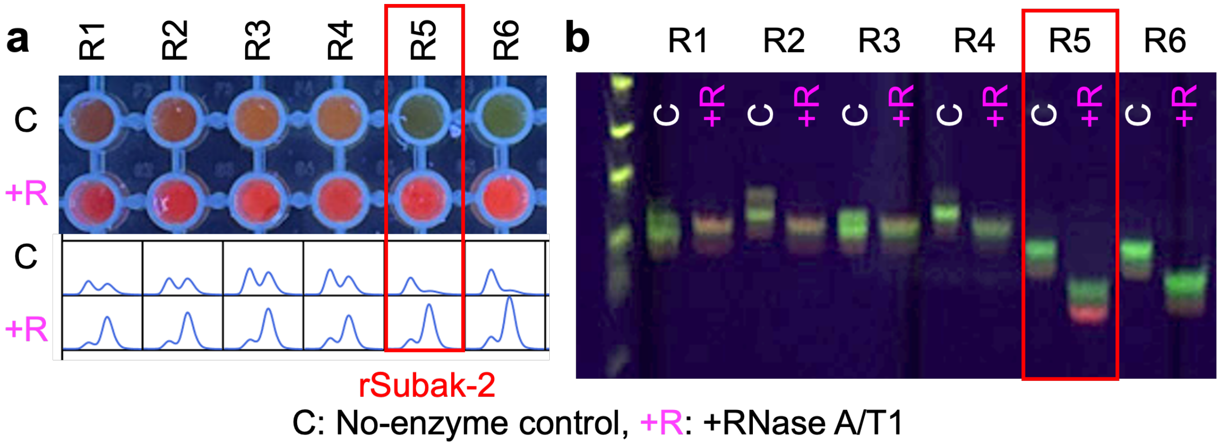

**Supplementary Fig. 14. Evaluation of Subak-2 variants for optimized Cas13a accessibility and background signal.** Fluorescence and electrophoretic characterization of Subak-2 variants (R1-R6) featuring various bulge motifs (R1-R4) and ribonucleotide substitutions (R5, R6). (**a**) Fluorescence imaging in microplates (top) alongside intensity profiles (bottom) compare intact (No-enzyme control) and cleaved (+RNase A/T1) states. While the incorporation of bulge motifs improves enzymatic accessibility, a concomitant increase in pre-cleavage red emission is observed. (**b**) PAGE analysis reveals that the variant featuring six consecutive ribonucleotides (R5, red box) maintains minimal pre-cleavage red emission while yielding robust post-cleavage red, identifying it as the optimal construct for high signal-to-noise ratio. All reactions were performed with 10 µM substrate in SPB (pH 7.4) at 37 °C for 30 min. Sequence information is in **Supplementary Table 3**.

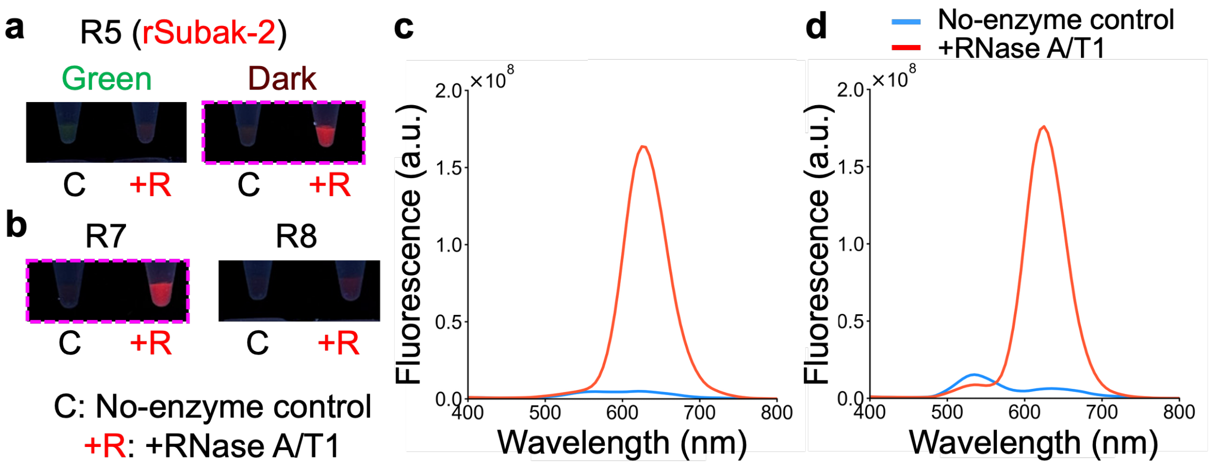

**Supplementary Fig. 15. Investigation of double-sided cleavage for improved signal-to-noise ratio.** To evaluate whether double-sided cleavage enhances red emission, additional ribonucleotide substitutions were introduced to the right stem of the R5 (rSubak-2) scaffold. (**a**,**b**) Comparison of fluorescence emission between the original R5 and extended variants (R7, R8). Purification of the dark state variants (**a**, highlighted in magenta dashed box) ensures a complete transition to distinct red fluorescence upon RNase A/T1 digestion. (**c**, **d**) Fluorescence emission spectra of the optimized variants (c) R5 and (d) R7, corresponding to the magenta dashed boxes in (a) and (b), respectively. While additional cleavage sites produce a robust red signal, certain variants exhibit a post-cleavage green fluorescence, indicating a trade-off between cleavage efficiency and background signal.

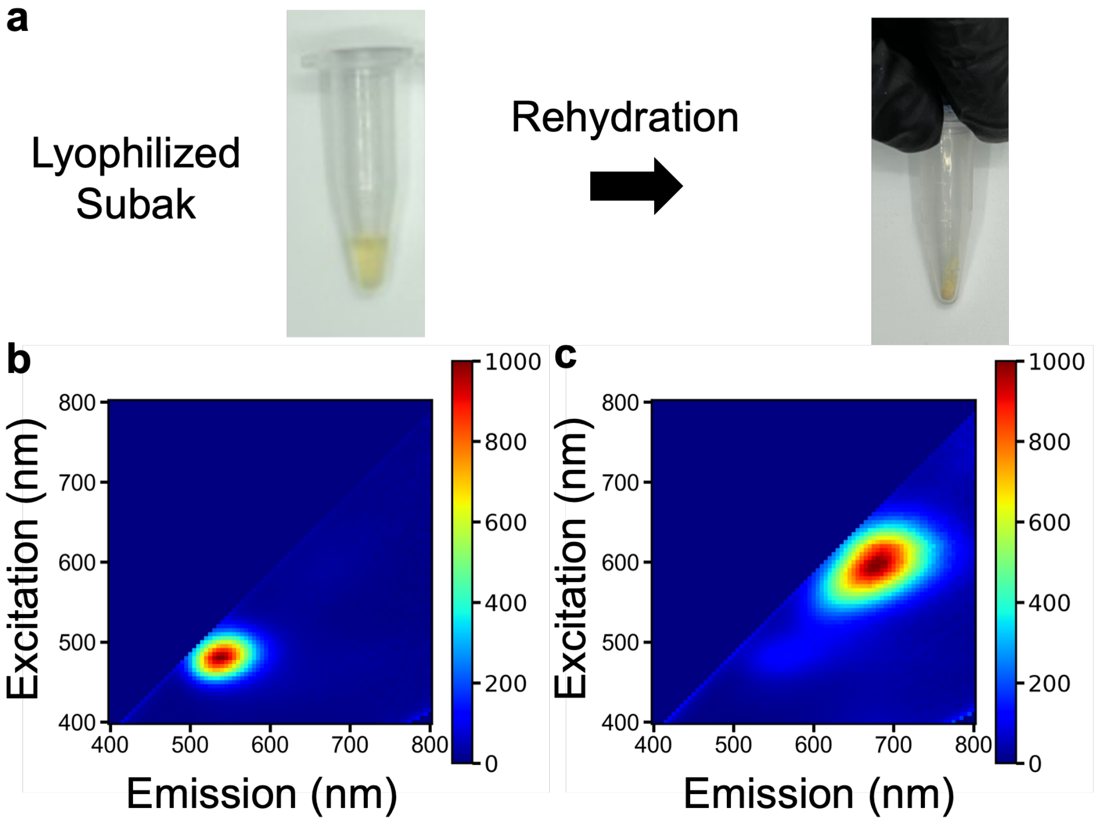

**Supplementary Fig. 16. Stability and performance of lyophilized Subak-1.** (**a**) Lyophilized Subak-1 retains its characteristic color transition upon rehydration. Excitation-emission matrix (EEM) spectra of the re-suspended Subak-1 (**b**) before and (**c**) after DNase I cleavage.
